# Detailed and atypical HLA-E peptide binding motifs revealed by a novel peptide exchange binding assay

**DOI:** 10.1101/2020.05.05.078790

**Authors:** Lucy C. Walters, Andrew J. McMichael, Geraldine M. Gillespie

## Abstract

Diverse SIV epitopes that bind the rhesus homologue of HLA-E, Mamu-E, have recently been identified in SIV-vaccine studies using a recombinant Rhesus cytomegalovirus (RhCMV 68-1) vector, where unprecedented protection against SIV challenge was achieved. Additionally, several *Mycobacterial* peptides identified both algorithmically and following elution from infected cells, are presented to CD8^+^ T cells by HLA-E in humans. Yet, a comparative and comprehensive analysis of relative HLA-E peptide binding strength via a reliable, high throughput *in vitro* assay is currently lacking. To address this we developed and optimised a novel, highly sensitive peptide exchange ELISA-based assay that relatively quantitates peptide binding to HLA-E. Using this approach we screened multiple peptides, including peptide panels derived from HIV, SIV and Mtb predicted to bind HLA-E. Our results indicate that although HLA-E preferentially accommodates canonical MHC class I leader peptides, many non-canonical, sequence diverse, pathogen-derived peptides also bind HLA-E, albeit generally with lower relative binding strength. Additionally, our screens demonstrate that the majority of peptides tested, including some key Mtb and SIV epitopes which have been shown to elicit strong Mamu-E-restricted T cell responses, either bind HLA-E extremely weakly or give signals that are indistinguishable from the negative, peptide-free controls.

## Introduction

It is well established that HLA-E preferentially accommodates a signal peptide comprising residues 3-11 of MHC class Ia leader sequences, typically VMAPRTL(V/L/F)L (VL9), in its binding groove and that these peptides dominate the HLA-E-presented ligandome in the steady state ^1 2^. VL9 peptide-bound HLA-E complexes constitute major ligands for heterodimeric inhibitory CD94-NKG2A and activating CD94-NKG2C receptors predominantly expressed on NK cells. VL9-bound HLA-E-CD94/NKG2A engagement has been shown to regulate NK cell-mediated lysis and represents an important component of immune homeostasis ^3 4 5^.

In the original peptide binding studies, the HLA-E sequence motif was explored via Ala and Gly substitution experiments along the HLA-B*0801 leader peptide (VMAPRTVLL), for which strong sequence selectivity was uncovered at primary anchor positions 2 (Met) and 9 (Leu), with weaker preferences observed for secondary anchor positions 7 (Val) and 3 (Ala) ^1^. Subsequent structural characterisation of HLA-E revealed unusually high binding pocket occupancy compared to classical MHC class Ia molecules. In addition to the peptide’s primary anchor side chains at positions 2 (Met) and 9 (Leu) that project into their respective B and F pockets, positions 3 (Ala), 6 (Thr) and 7 (Val/Leu) also constituted secondary anchor residues, with their side chains occupying the shallow D, C and deeper E pockets, respectively ^6 7 8^. The shape complementarity of these side chains, in addition to pocket-based interactions such as hydrogen bonding, van der Waals and hydrophobic contacts place sequence restrictions on the peptide in these various anchor positions, reducing the overall magnitude of the potential peptide binding repertoire ^9^. It was argued therefore that the requirement to occupy a greater number of pockets, as observed in VL9-bound structures of HLA-E, would correlate with a less diverse peptidome than classical MHC class I allotypes ^6^. However, non-VL9 sequence-diverse peptides that bind HLA-E or its rhesus or murine orthologues, Mamu-E or Qa-1 respectively, have subsequently been identified in diverse immunological settings. For example, HLA-E- and Qa-1-restricted peptides from pathogens including Salmonella ^10,11^, Hepatitis C virus ^12^, Epstein-Barr virus (EBV) ^13^ and Influenza virus ^13^, have been reported. These findings also extend to self-derived peptides from proteins such as Fam49B ^14^, Hsp60 ^15^ and Prdx5 ^16^ in the context of defective antigen processing, cellular stress or autoimmune disease.

More recently, two contexts where large numbers of MHC-E-restricted epitopes are presented have been reported - vaccination of rhesus macaques with a RhCMV 68-1 vector recombinant for SIV or HIV antigen ^17^ and *Mycobacteria tuberculosis* (Mtb) infection, BCG vaccination or environmental *Mycobacterial* exposure ^18,19^. Importantly, Mamu-E-restricted CD8^+^ T cell responses have been implicated as immune correlates of unprecedented protection against SIV challenge in RhCMV68-1 vector vaccination studies in rhesus macaques ^17^. However, these Mamu-E-restricted epitopes are sequence-diverse, contain non-canonical anchor residues and lack a common motif ^17^. Notably, despite a considerable degree of sequence divergence between the human and rhesus MHC-E homologues, heavy chain-derived residues forming the B, D, C, E and F pockets of the binding groove exhibit strong sequence conservation ^20^. Given this stringent sequence conservation between pocket-forming residues, it is likely these homologues share a largely overlapping peptide binding repertoire. In the second predominant setting, 69 *Mycobacterial* peptides were predicted to bind HLA-E *in silico* using peptide motif-based algorithms, of which 55 were shown to stimulate HLA-E-restricted CD8^+^ T cell responses in PPD-responding individuals ^18^. A smaller set of HLA-E-restricted Mtb-derived peptides were more recently eluted from infected cells and validated as CD8^+^ T cell epitopes ^21^.

Given the identification of a large number of non-VL9 pathogen- and self-derived HLA-E-binding peptides, we recognised the need to develop an assay to reliably and comparatively assess the HLA-E peptidome. We therefore designed and optimised a new *in vitro* peptide exchange ELISA-based assay to validate and relatively quantitate peptide binding to HLA-E. Although we previously validated HLA-E peptide binding^17^ using a micro-refolding ELISA-based assay, originally developed to determine peptide binding affinity to classical MHC class I molecules ^7 22^, this assay suffered from variability between biological repeats, so that very large numbers of replicate assays were required to achieve statistically robust results. The variability was likely caused by assay-to-assay fluctuations in protein refolding in addition to unusually high background signals, due to partial HLA-E-β2M refolding in the absence of added peptide ^20^. To improve the system we adopted a new UV peptide exchange ELISA-based assay ^23^. In this assay, HLA-E heavy chain and β2M are pre-refolded with a UV-labile HLA-E binding peptide based on the VL9 sequence in which the solvent exposed Arg at position 5 of VL9 was replaced with a photo-cleavable, unnatural 2-nitrophenylamino acid residue ^20 23^. Such conditional peptide ligand technology was originally developed for HLA-A*02:01 ^23^ and subsequently adapted to facilitate peptide exchange in HLA-A*1, -A*3, -A*11, -B*7 and -B*57 molecules ^24 25^. Specifically, the nitrophenyl moiety within the substituted beta amino acid is sensitive to UV illumination at >350nm. This results in peptide cleavage and successive dissociation of the degraded peptide fragments from the MHC class I peptide binding groove, as previously validated by mass spectrometric analysis ^23^. Consequently, in the absence of bound peptide, MHC class I complexes become destabilised and disintegrate. However, complex disintegration can be circumvented by peptide exchange with a rescue peptide that binds the MHC allele in question.

Using this approach both pathogen- and self-derived peptides previously reported to bind HLA-E were screened. Our results indicate that although HLA-E preferentially accommodates MHC class I leader peptides, a range of pathogen-derived peptides containing hydrophobic and polar primary anchor residues are also tolerated, albeit with lower relative binding strength relative to VL9 peptides. Importantly, our screens also demonstrate that the majority of peptides tested, including some key Mtb and SIV epitopes which have been shown to elicit immunodominant MHC-E-restricted T cell responses, bind HLA-E extremely weakly or generate indistinguishable signals from the no peptide negative controls. Finally, while the stronger binding peptides show a clear sequence motif that could help identify further antigenic peptides, many of the well-defined peptide epitopes lack a motif suggesting that alternative peptide binding modes may exist for these peptides *in vivo*.

## Results

### Micro-refolding versus UV peptide exchange HLA-E binding assay comparison

We conducted a direct comparison of the previously published HLA-E micro-refolding ^20 7^ versus UV-assisted peptide exchange binding assays to determine which approach yielded the highest sensitivity and reliability in discriminating weak HLA-E peptide binders from background signals (Figure 1). Three biological repeats for two test peptides, BZLF1 (SQAPLPCVL) and GroEL (KMLRGVNVL) which were previously reported to support HLA-E or Qa-1 stabilisation, were included in this comparison ^13 10^. Notably, the hierarchy of relative peptide affinity remained constant across both approaches with the BZLF peptide exhibiting the strongest and GroEL the weakest binding to HLA-E (Figure 1, A). However, the average absorbance reading for each test peptide was statistically significant in the initial UV-induced peptide exchange assay, whereas only VL9 achieved significance in the micro-refold approach. Further, the standard deviation between biological repeats was lower for all peptides tested including the positive and negative controls in the UV-induced peptide exchange assay compared to the micro-refold approach (Figure 1, B). The standard deviation for three biological repeats of the ‘no peptide’ negative control decreased by over an order of magnitude in the UV exchange approach compared to the micro-refold assay (from 0.49 to 0.02, respectively).

**Figure 1.**
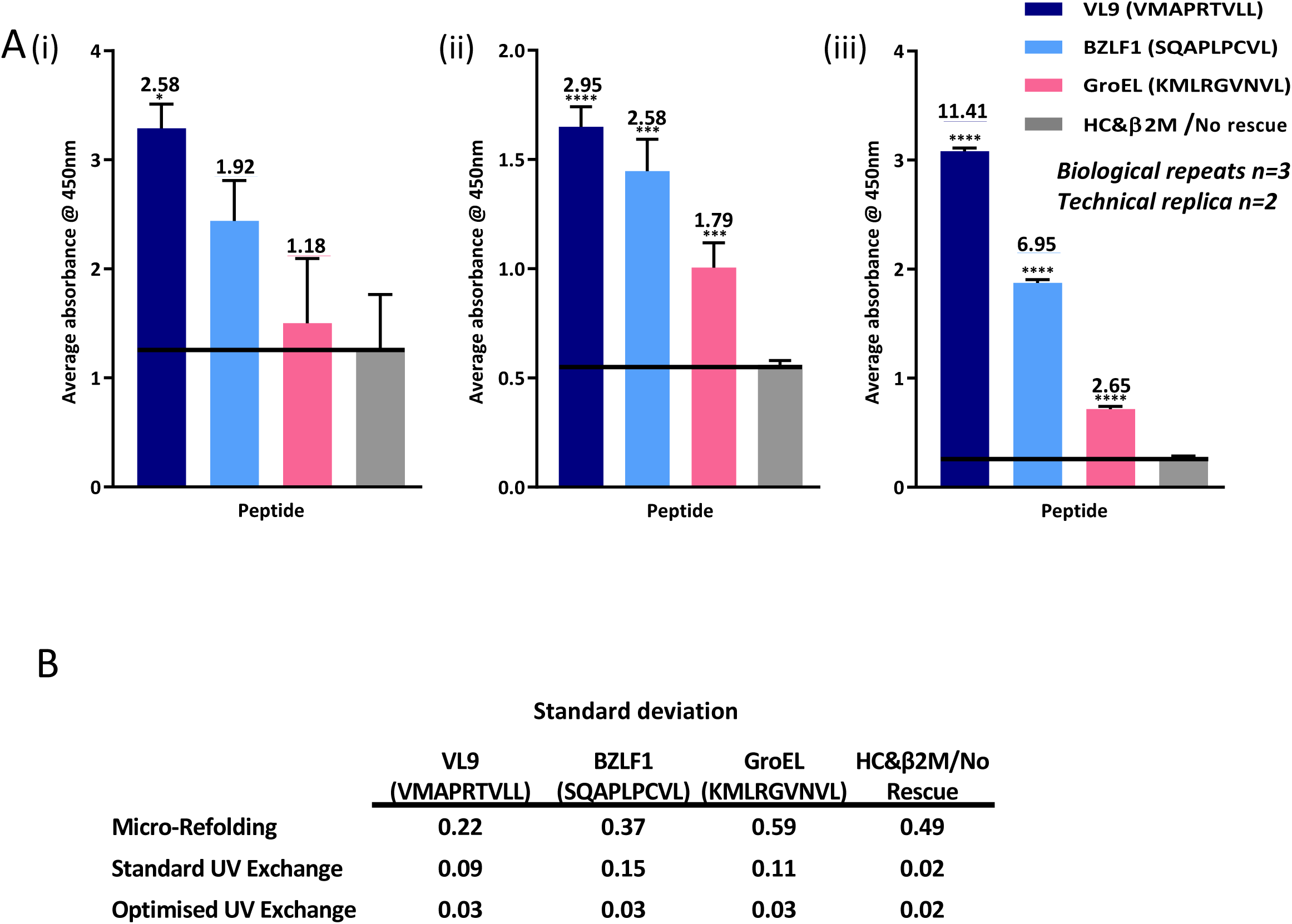
Comparison of micro-refold versus UV exchange peptide binding assay approaches. **A.** Average absorbance readings for 2 HLA-E binding pathogen-derived peptides sampled in **(i)** the previously published micro-refolding peptide binding affinity assay, **(ii)** the original (standard) UV-exchange peptide binding affinity approach and **(iii)** the optimised UV exchange peptide binding assay approach. The Y-axis denotes average absorbance readings at 450nm and the X-axis denotes test peptides, which are colour-coded according to the shared Figure legend. Peptide to background ratios are indicated above each bar, with peptide ID and sequences denoted. The sequence origin and corresponding references are detailed in Supplementary Table 1, C. *The mouse anti-human HLA-E monoclonal antibody 3D12 was used as the ELISA capture antibody, with polyclonal rabbit anti-human β2M HRP-conjugated IgG antibody used as detection, followed by goat-anti-rabbit EnVision+ System-HRP signal amplification. The positive control peptide VL9 (HLA-B7-derived leader sequence peptide, VMAPRTVLL) is shown in dark blue. Peptide free negative background are noted in grey; for micro-refold approach this consisted of HLA-E heavy chain and β2M only (HC&β2M) whereas the UV peptide exchange approach negative control comprised previously refolded HLA-E complexes with no added rescue peptide in the exchange reaction (No Rescue). Three biological repeats (three individual peptide exchange reactions per test peptide) were carried out simultaneously, with two technical replicas sampled from the same peptide exchange reaction performed for each biological repeat. (reported herein as n=3, technical replica n=2). Stars above error bars reflect degree of significance in peptide binding by two-tailed t-tests: no star=P>0.05, *=P≤0.05, **=P≤0.01, ***=P≤0.001, ****P=≤0.0001.* Error bars depicting the mean and standard deviation of biological repeats are shown. *The conditions outlined in italics are replicated in all subsequent ELISA optimisation assays.* **B.** Table of standard deviation for the micro-refold and the standard and optimised UV exchange peptide binding assay approaches.

### UV peptide exchange HLA-E binding assay optimisation

#### a. Optimisation of UV peptide exchange ELISA-based assay buffer system

Three distinct buffer systems were trialled in the photo-assisted peptide exchange reaction including a Lutrol-Tris-maleate-based buffer previously used in the micro-refolding peptide binding assay ^22 7 20^, a Tris-based buffer previously used for UV-induced peptide exchange with classical MHC class I molecules ^23^, and finally an L-Arginine-Tris-based buffer traditionally used in macro-refolding of class I MHC proteins ^26^ (Figure 2, A, i). The L-Arginine-Tris-based buffer yielded lower peptide-free background signals, concomitantly improving peptide to background ratios for all peptides tested. As the peptide to background ratios were superior when the peptide exchange reaction was conducted in the L-Arginine-Tris-based buffer, this buffer was subsequently adopted for follow-up. To determine which component of the L-Arginine-Tris-based buffer was responsible for reduced peptide-free background signals, further assays were carried out in which L-Arginine monohydrochloride or the glutathione redox components were individually excluded (Figure 2, A, ii). In the absence of L-Arginine monohydrochloride, the average absorbance signal for the peptide-free background increased from 0.18 to 0.65, whereas, in the absence of the redox system the background signal remained unaffected at 0.18. Such results indicate that L-Arginine monohydrochloride is the reagent responsible for peptide-free background reduction. To further evaluate the effect on the background signal, the concentration of L-Arginine monohydrochloride was titrated and an inverse correlation was observed between L-Arginine monohydrochloride concentration and the average absorbance reading of the peptide-free negative control (Figure 2, A, iii). As lower peptide to background ratios, including those for the positive control VL9 peptide, were also observed at increasing L-Arginine monohydrochloride concentrations, 400mM was chosen as the optimal concentration for subsequent assays.

**Figure 2.**
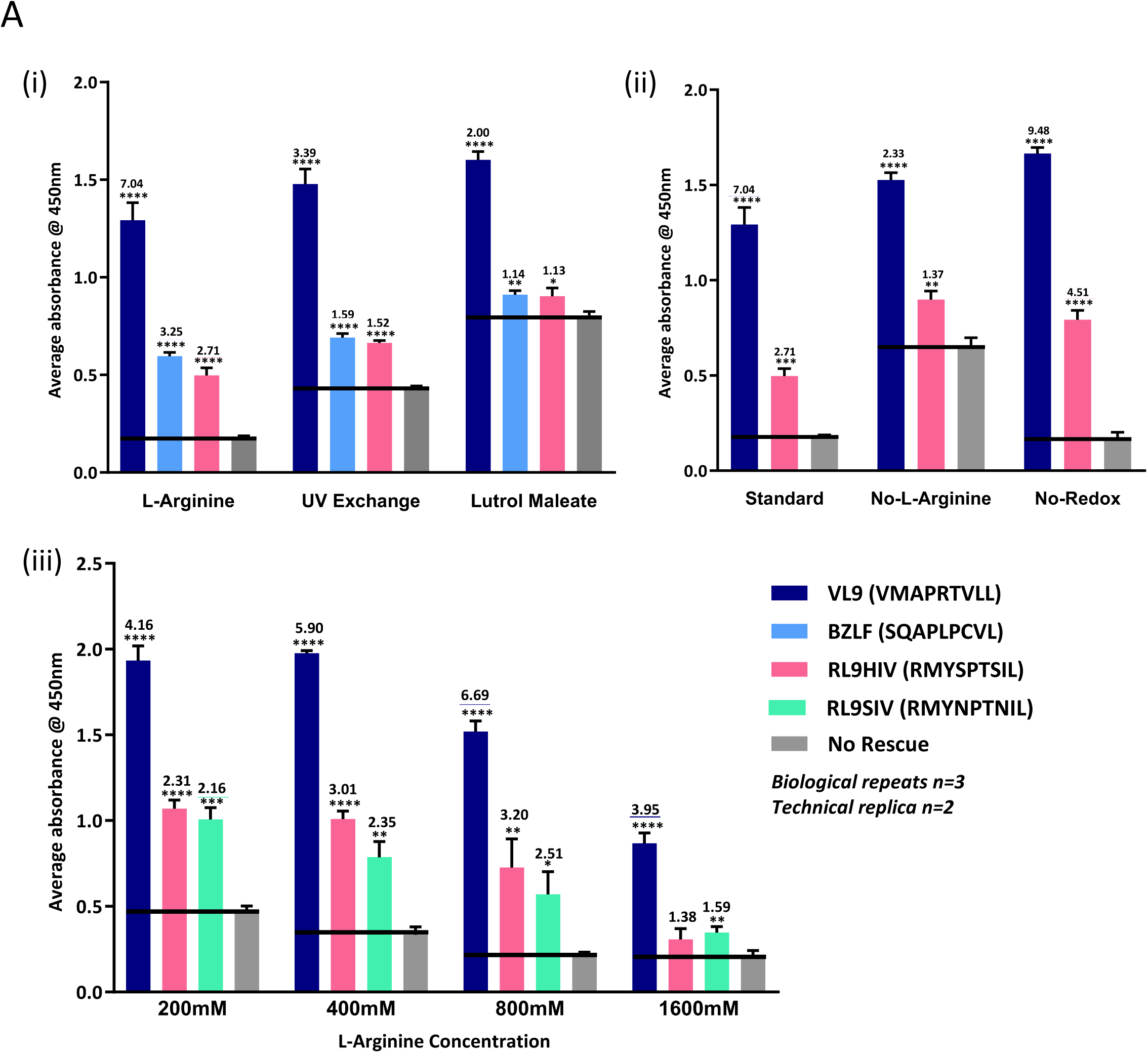
UV-induced peptide exchange buffer optimisation. **A. (i)** Average absorbance readings from 3 different buffer systems trialled in the UV-induced peptide exchange sandwich ELISA peptide binding assay. The Y-axis denotes average absorbance readings at 450nm and the X-axis denotes the buffer system. Peptide to background ratios are indicated above each bar. **(ii)** Average absorbance readings from 3 peptide binding ELISA optimisation assays in which components of the L-Arginine-based refolding buffer were individually excluded. The ‘standard’ buffer contains all components of the L-Arginine-based macro-refolding buffer (detailed in Materials and Methods). L-Arginine monohydrochloride was excluded from the ‘No L-Arginine’ assay and the glutathione redox system excluded from the ‘No-Redox’ assay. The Y-axis denotes average absorbance readings at 450nm and the X-axis denotes buffer composition. Peptide to background ratios are indicated above each bar. **(iii)** Average absorbance readings from peptide binding assays in which L-Arginine monohydrochloride concentration was varied in the UV-induced peptide exchange reaction from 200mM to 1600mM. The Y-axis denotes average absorbance readings at 450nm and the X-axis denotes L-Arginine concentration. Peptide to background ratios are indicated above each bar. Shared Figure legend colouring throughout, with individual peptide IDs and sequences denoted. Peptide ID, sequence, origin and corresponding references are detailed in Supplementary Table 1, A (iv) and C. Error bars depicting the mean and standard deviation of biological repeats are shown.

#### b. UV exposure versus no-exposure optimisation

The recommended duration of UV exposure in previously published protocols using UV-labile ligand technology is 1hr ^23^. However, we hypothesised that extended exposure could enable further cleavage of UV-sensitive peptides resulting in enhanced destabilisation of HLA-E complexes in the absence of rescue peptide that should, in turn, further reduce background signals and improve peptide to background ratios. We therefore exposed the peptide exchange reaction to UV radiation for zero, one or two hours before sampling by sandwich ELISA (Figure 3, A, i). As predicted, the peptide to no rescue background ratios of both test peptides, RL9HIV (RMYSPTSIL) and RL9SIV (RMYNPTNIL), and the positive control VL9 peptide were higher after two hours of UV exposure. Further investigation revealed that peptide to background ratios of both test peptides and the positive control VL9 were similar after two or three hours of UV exposure and began to decrease at the four hour UV exposure time-point, perhaps due to radiation damage reducing overall protein yield in the reaction (Figure 3, A, ii). However, it remained uncertain whether this increased assay sensitivity was attributable to prolonged photo-irradiation or simply due to the extended time period over which peptide exchange could occur. Although exposure to light within the UV spectrum was originally deemed necessary for photo-cleavage and resultant exchange of HLA-A2.1-restricted UV-labile peptides ^23^, this had not been fully elucidated for the UV-labile HLA-E binding peptide (VMAPJTLVL) ^20^. Also, since we previously observed unassisted peptide exchange in the blue native gel system where pre-refolded VL9-bound HLA-E incubated in the presence of excess Mtb44 was capable of converting to Mtb44 peptide-loaded material ^20^ (evidenced by distinct Blue Native gel band positioning), experiments to test the possibility of UV-independent peptide exchange were therefore conducted (Figure 3, A, iii). Both in the presence or absence of UV irradiation we observed similar data trends for test peptide to background ratios over time-points of zero to three hours indicating peptide exchange did not require targeted UV illumination at 366nm. However, in contrast to photo-illuminated peptide exchange reactions, test peptide to background ratios increased beyond the three hour time-point in the absence of UV. An optimal duration of 5 hours was thus selected as standard for the non-UV peptide exchange reaction.

**Figure 3.**
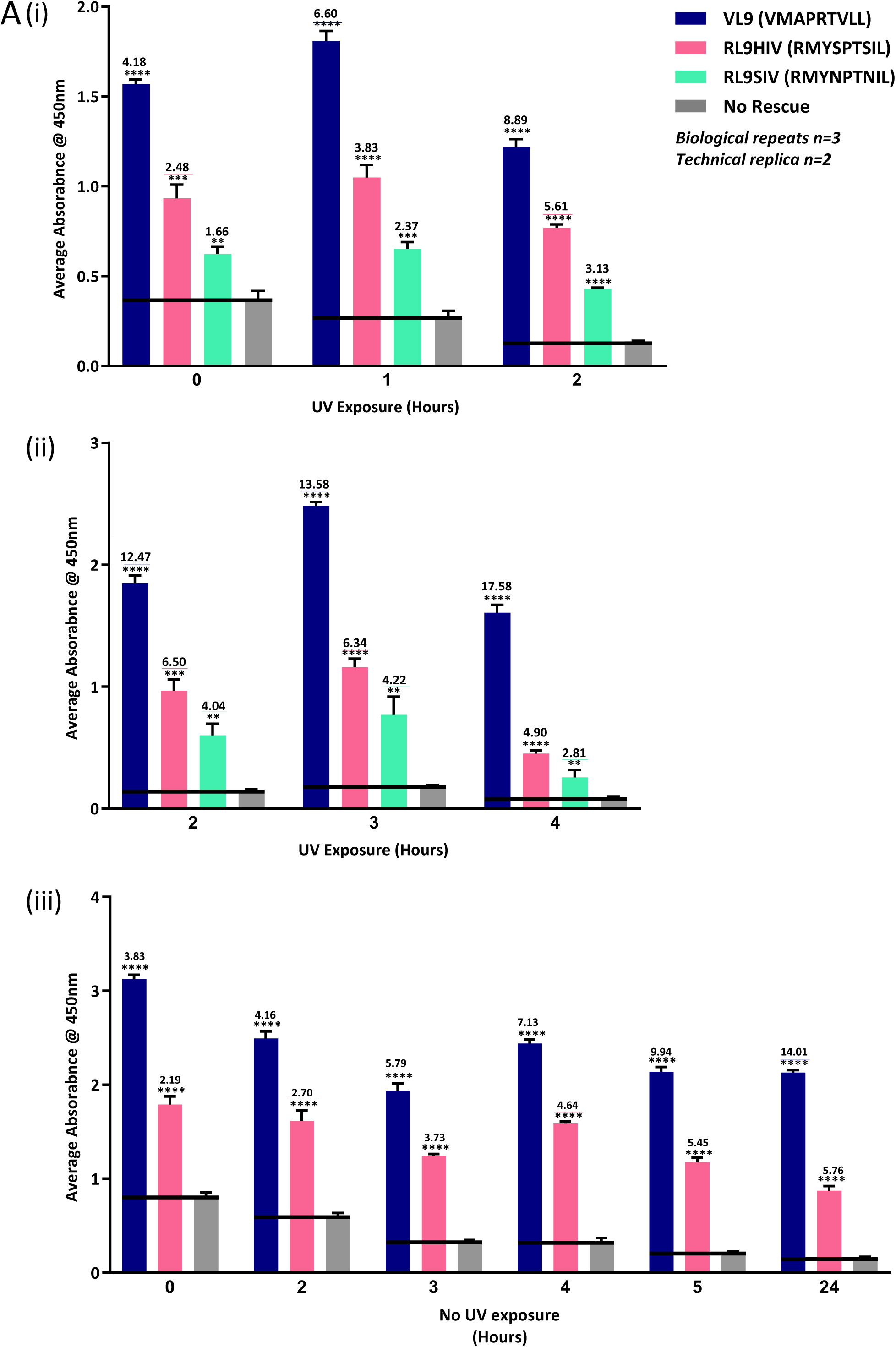
Optimisation of UV exposure duration in the peptide exchange reaction. **A. (i)** Average absorbance readings from three sandwich ELISAs where the peptide exchange reaction was sampled at 0, 1, and 2 hours of UV exposure. The Y-axis denotes average absorbance readings at 450nm and the X-axis denotes duration of UV exposure. Peptide to background signal ratios are denoted above error bars. **(ii)** Average absorbance readings from 3 sandwich ELISAs where the peptide exchange reaction was sampled at 2, 3, and 4 hours of UV exposure. The Y-axis denotes average absorbance readings at 450nm and the X-axis denotes duration of UV exposure. Peptide to background signal ratios are denoted above error bars. **(iii)** Average absorbance readings from 6 sandwich ELISAs where the peptide exchange reaction was sampled at 0, 2, 3, 4, 5, and 24 hours in the absence of UV exposure. The Y-axis denotes average absorbance readings at 450nm and the X-axis denotes duration of the peptide exchange reaction at 4°C. Peptide to background signal ratios are denoted above error bars. Shared Figure legend colourings are denoted with individual peptide IDs and sequences listed. Peptide ID, sequence, origin and corresponding references are detailed in Supplementary Table 1, A (iv). Error bars depicting the mean and standard deviation of biological repeats are shown.

Following these optimisations, greater statistical significance was achieved for both BZLF1 (SQAPLPCVL) and GroEL (KMLRGVNVL) test peptides and the standard deviation was further reduced relative to the non-optimised UV peptide exchange assay, except for the negative control which remained low at 0.02 (Figure 1). Assay sensitivity, defined by an ability to discriminate between low affinity HLA-E peptide binding and the no-peptide background, was considerably improved post-optimisation – the test peptide to background signal ratio for BZLF1 more than doubled relative to the initial UV exchange assay.

### Screening of HIV-derived peptides predicted to bind HLA-E

Once optimisation protocols for the peptide exchange ELISA-based assay were complete, peptide panels comprising HIV-derived 9mer peptides were screened for their HLA-E binding capacity. The first panel consisted of peptides identified in-house as HLA-E predicted binders using the NetMHC 4.0 web-based server ^27,28^ (Figure 4, A), and thus were biased by computational predications based on previously established HLA-E binding motifs. 11 of the predicted peptides had the canonical Leu at position 9 and 11 contained a hydrophobic residue at position 2. A range of relative binding affinities were observed for this panel, with 12 of the 16 peptides demonstrating significant binding to HLA-E, albeit with much lower strength compared to the VL9 control peptide. For example, five of the significant HLA-E binding peptides in this panel generated average absorbance readings below 5% of the positive control VL9 signal following background subtraction (Supplementary Table 2). A second panel comprising HIV Gag- and Pol-derived peptides, based on a simple prediction of canonical primary anchor position 2 Met and a hydrophobic primary anchor position 9 residue, were also screened (Figure 4, B). Although two peptides from this panel demonstrated significant binding above background, all peptides exhibited very low average absorbance signals, only slightly higher than the negative control peptide-free background and all below 5% of the positive control VL9 signal following buffer subtraction (Supplementary Table 2). Peptides containing overlapping HIV Gag 9mers comprised the third panel screened for HLA-E binding. Many of these peptides contained biochemically distinct, non-canonical primary and secondary anchor residues and would not qualify as HLA-E binding peptides based on previously established sequence motif algorithms. The majority of these peptides generated exceptionally low binding signals with average absorbance readings only marginally above the no rescue background signal. The relative HLA-E peptide binding strength of a small peptide panel comprising the immunodominant SIV-derived RL9SIV (RMYNPTNIL: termed ‘supertope’ because it elicits CD8^+^ T cell responses in all rhesus monkeys vaccinated by the RhCMV 68-1 vaccine^17^) and its HIV Clade B-derived alignment counterpart peptide, RMYSPTSIL (peptide ID: RL9HIV), was also evaluated using the peptide-exchange ELISA-based assay (Figure 4, D). The physiological relevance of the RL9HIV peptide was validated by the identification of specific Mamu-E-restricted CD8^+^ T cell responses in rhesus macaques vaccinated with RhCMV 68-1 vectors recombinant for HIV Gag ^20^. Although both RL9HIV and RL9SIV binding exhibited the greatest degree of statistical significance (*P=≤0.0001)*, the average absorbance signal for RL9HIV was consistently greater than that generated by RL9SIV; the RL9HIV average absorbance reading comprised 43% of the positive control VL9 signal, whereas RL9SIV was lower at 22% (Supplementary Table 2).

**Figure 4.**
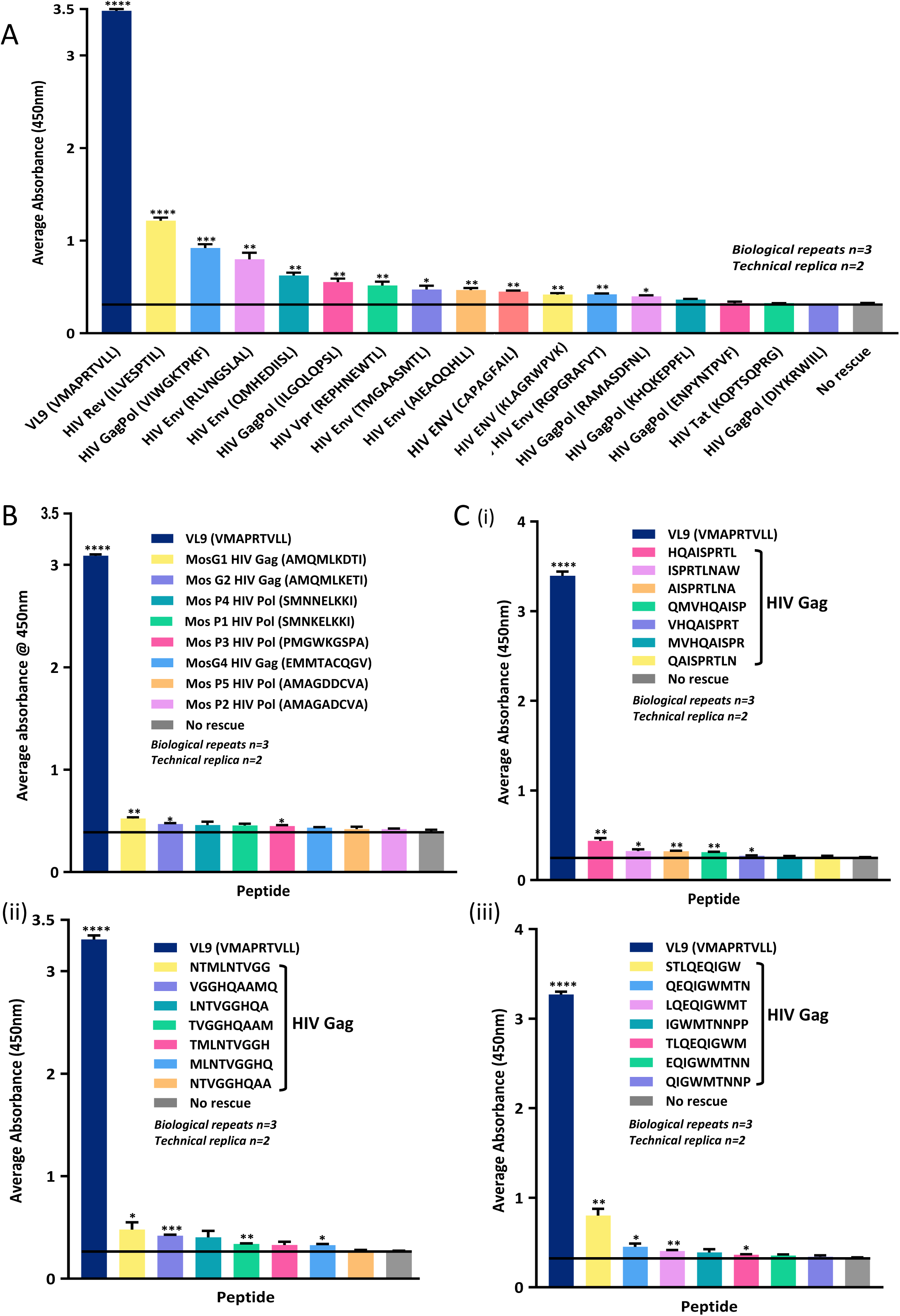
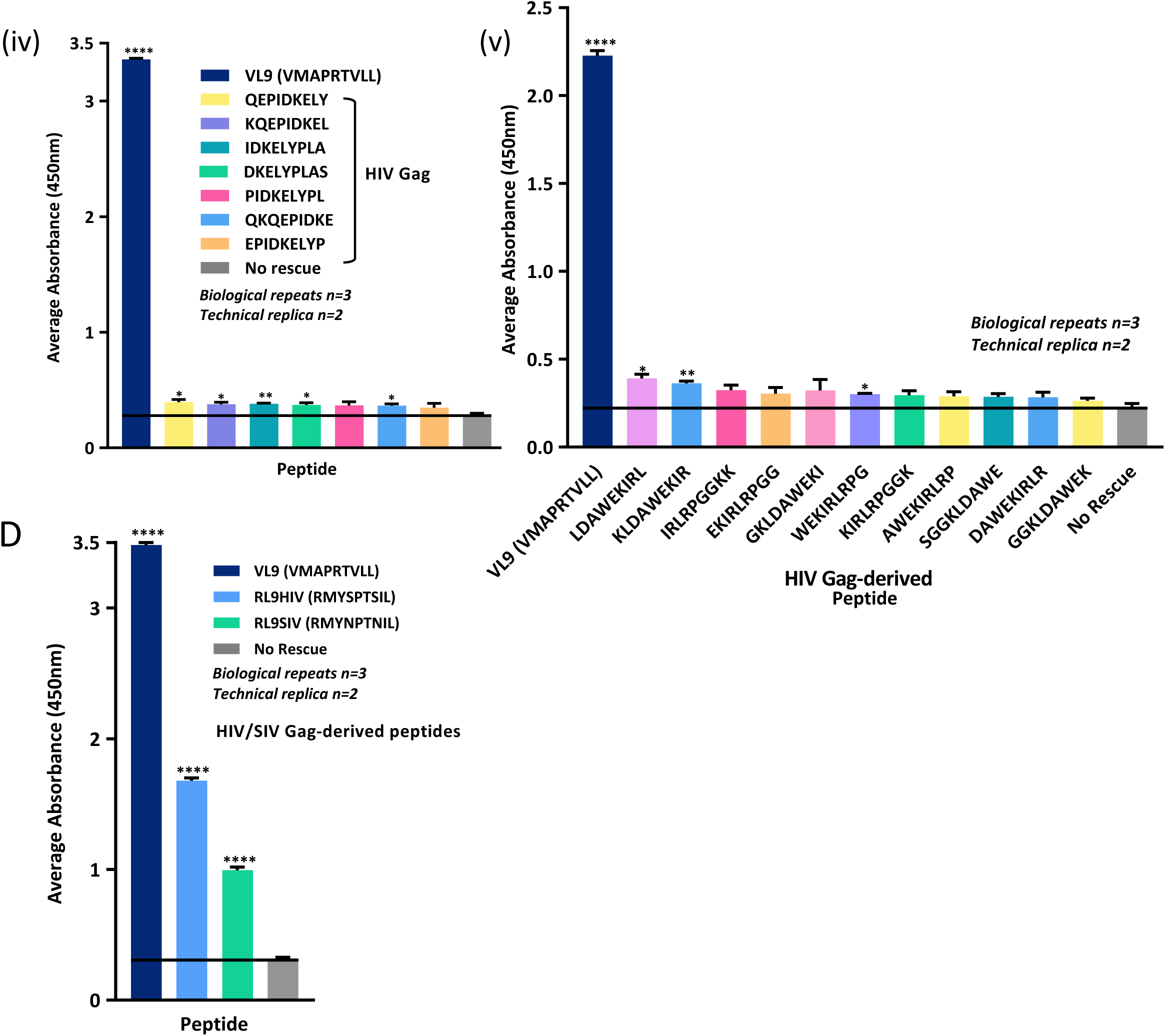
Screening of HIV-derived peptides in the HLA-E peptide exchange ELISA assay. **A.** Average absorbance readings obtained in the peptide exchange ELISA-based assay for HIV-derived peptides predicted to bind HLA-E by NetMHC. Peptides tested in the peptide exchange-based binding assay are denoted on the X-axis, with average absorbance readings at 450nm on the Y-axis. Peptide ID, sequence, origin and corresponding references are detailed in Supplementary Table 1, A (i). **B.** Average absorbance readings for HIV Gag and Pol derived peptides which contain the canonical position 2 primary anchor residue (Met) and a hydrophobic position 9 primary anchor residue. Peptide ID, sequence, origin and corresponding references are detailed in Supplementary Table 1, A (ii). **C. (i), (ii), (iii), (iv) & (v)** Average absorbance readings for overlapping 9mer peptides derived from HIV Gag that were screened for HLA-E binding capacity in the peptide exchange ELISA-based binding assay. Peptide ID, sequence and origin are detailed in Supplementary Table 1, A (iii). **D.** Average absorbance readings for RhCMV 68-1 vaccine-identified RL9SIV and RL9HIV peptide epitopes tested in the peptide exchange ELISA-based binding assay. Peptide ID, sequence, origin and corresponding references are detailed in Supplementary Table 1, A (iv). Average absorbance readings at 450nm are displayed on the Y-axis for plots in **B, C** and **D**. Error bars depicting the mean and standard error of the mean (SEM) of biological repeats are shown.

### Screening of Mtb-derived peptides predicted to bind HLA-E

A panel of *Mycobacterial* peptides eluted from Mtb-infected cells ^21^ was screened in the improved assay system (Figure 5, A). As these peptides were eluted from HLA-E, they were not subject to the algorithmic biases that dictate previously described HLA-E-restricted peptide sequence motifs. Peptides in this panel not only deviated from the canonical HLA-E sequence motif but also from the preferred nonameric peptide length. In addition to a peptide comprising 12 amino acids (GT12, GGILIGSDTDLT), two nonameric Mtb peptides, IL9 (IMYNYPAML) and LL9 (LLDAHIPQL), with hydrophobic residues at both primary anchor positions demonstrated significant binding to HLA-E. The IL9 signal was particularly strong and represented 65% of the VL9 positive control signal following background subtraction (Supplementary Table 2). A small subset of the 69 previously published *Mycobacterial* peptides that were algorithmically predicted to bind HLA-E ^18^, were also screened (Figure 5, B). As these peptides were initially identified *in silico* based on previous sequence motifs, they are largely canonical with either a hydrophobic Leu or Met at position 2 and a hydrophobic Leu or Ile at position 9. One peptide, Mtb44 (RLPAKAPLL), exhibited strikingly strong binding, generating an average absorbance signal that comprised 97% of the VL9 positive control signal following buffer subtraction (Supplementary Table 2) and represented the only non-VL9 peptide screened in the ELISA assay to generate an average absorbance reading within the standard error of the mean of the positive VL9 control. The second strongest binder, Mtb14 (RMAATAQVL), generated average absorbance readings between 40-50% of the positive control signal following background subtraction and contained the canonical primary anchor residues Met at position 2 and Leu at position 9 (Supplementary Table 2). Although all peptides in this panel generated average absorbance signals above background levels in the peptide exchange assay, Mtb54 (FLLPRGLAI), Mtb48 (RLANLLPLI) and Mtb55 (VMATRRNVL) generated lower binding signals, despite Mtb55 containing canonical primary anchor residues.

**Figure 5.**
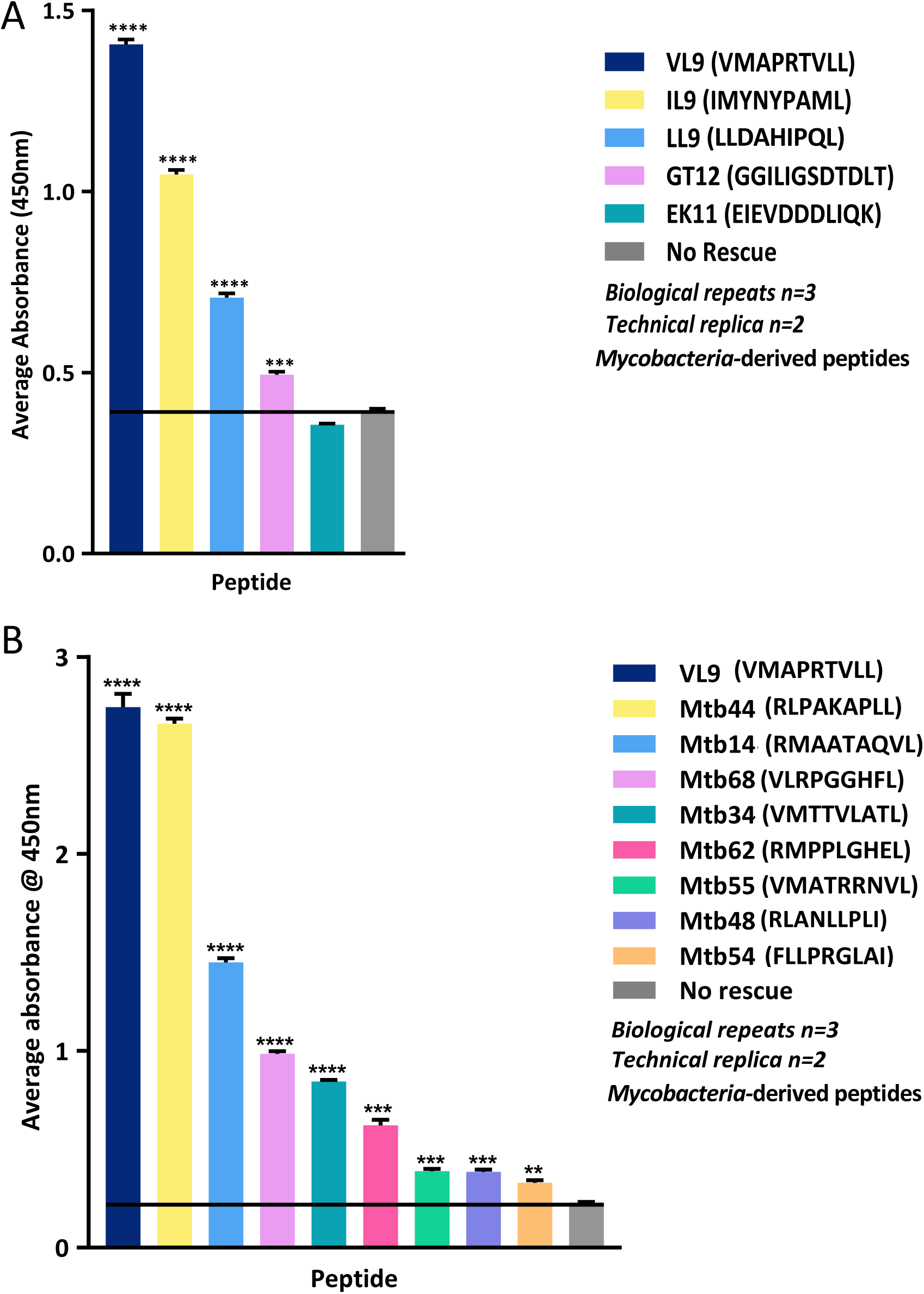
Screening of *Mycobacteria*-derived peptides in the HLA-E peptide exchange ELISA assay. **A.** Average absorbance readings from the peptide binding assay for Mtb-derived peptides eluted from infected cells. Peptide sequences are denoted in the corresponding Figure legend. Peptide ID, sequence, origin and corresponding references are detailed in Supplementary Table 1, B (i). **B.** Average absorbance readings from the peptide binding assay for Mtb-derived peptides previously predicted to bind HLA-E. Peptide sequences are denoted in the corresponding Figure legend. Peptide ID, sequence, origin and corresponding references are detailed in Supplementary Table 1, B (ii). Average absorbance readings at 450nm for plots in **A** and **B** are displayed on the Y-axis. Error bars depicting the mean and standard error of the mean (SEM) of biological repeats are shown.

### Screening of pathogen- and self-derived peptides predicted to bind HLA-E

A selection of previously described HLA-E-restricted peptides derived from a range of self-proteins or pathogens were screened in the ELISA assay ^10 12 13 29 14 15 30^ (Figure 6, A). A range of relative binding strengths was observed for these peptides including a Heat Shock Protein-derived peptide (Hsp5, GMKFDRGYI), a Fam49B peptide (FYAEATPML) and a Hep C Core protein-derived 10mer (YLLPRRGPRL) that generated average absorbance readings only marginally above that of the peptide-free control (Figure 6, A, i). Further, an Influenza Matrix peptide (ILGFVFTLT) previously reported to bind HLA-E yielded ELISA signals below that of the negative control. However, three peptides in this panel, including the EBV-derived BZLF1 (SQAPLPCVL), the Heat Shock Protein-derived Hsp4 (QMRPVSRVL) and the Salmonella-derived GroEL (KMLRGVNVL), produced ELISA signals greater than double the background signal. The Hsp4 and BZLF peptides generated particularly strong signals which respectively comprised 45% and 41% of the VL9 positive control signal following background subtraction (Supplementary Table 2). Although both GroEL and Hsp4 peptides contained the canonical primary anchor residues, position 2 Met and position 9 Leu, the BZLF1 peptide contained a polar Gln at position 2 in addition to a Cys in place of the canonical Val or Leu at the secondary anchor position 7. Further, despite the presence of canonical primary anchor residues, the Hsp4 peptide contains a charged Arg at both positions 3 and 7 in place of the preferred smaller canonical Ala at secondary anchor position 3 and the canonical hydrophobic Val or Leu at secondary anchor position 7 in VL9 peptides. Only one peptide in this panel, the Hepatitis C virus Core protein-derived peptide (YLLPRRGPRL), comprised a decamer sequence ^12^. However, this 10mer contains an internal nonameric sequence (LLPRRGPRL) which aligns more optimally to the canonical binding motif of HLA-E, with hydrophobic residues at positions 2 and 9. We thus compared binding of the LLPRRGPRL nonamer to the original decamer peptide and found that the average absorbance reading for the nonamer was more than double that of the original 10mer indicating that this shorter peptide is perhaps the optimal binder (Figure 6, A, ii).

**Figure 6.**
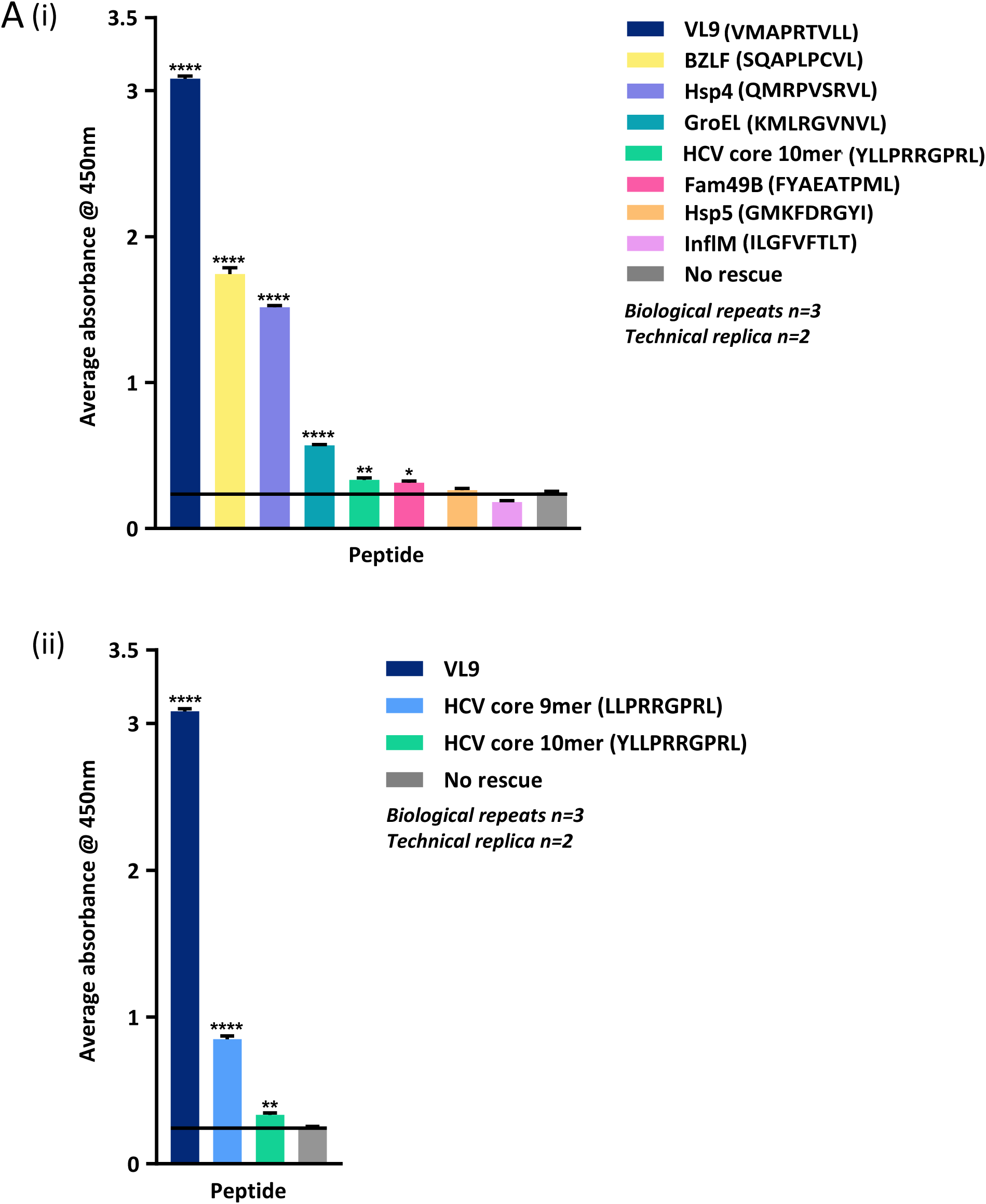
Screening of self- and pathogen-derived peptides previously predicted to bind HLA-E in the peptide exchange ELISA assay. **A. (i)** Average absorbance readings for pathogen- and self-derived HLA-E binding peptides screened in the peptide exchange ELISA-based assay for HLA-E binding capacity. Peptide sequences are denoted in the corresponding Figure legend. Peptide ID, sequence, origin and corresponding references are detailed in Supplementary Table 1, C (i). **(ii)** Peptide exchange ELISA-based assay comparison of the previously described Hepatitis C virus Core protein-derived decamer peptide and an internal nonameric predicted binding variant that adheres more optimally to the HLA-E binding motif and lacks the N-terminal Tyr of the parental peptide. Average absorbance readings at 450nm are displayed on the Y-axis for **(i)** and **(ii).** Error bars depicting the mean and standard error of the mean (SEM) of biological repeats are shown.

### Screening of high affinity HLA-E peptides with anchor residue mutations

To investigate HLA-E B pocket tolerability, VL9 peptide variants with substitutions at position 2 were synthesised and screened in the optimised peptide-exchange binding assay (Figure 7). First, a small panel based on the HLA-B7-derived leader peptide, VMAPRTVLL, was tested. Three of the four peptides in this panel, including a positively charged Lys and a polar Gln, demonstrated strong HLA-E binding, and generated average absorbance readings with the greatest degree of statistical significance (Figure 7, A, i). A second panel of position 2 anchor residue variants based on an alternative VL9 peptide (VMAPRTLLL) derived from the HLA-A1 leader sequence was also screened (Figure 7, A, ii). All 7 peptides in this panel demonstrated significant binding to HLA-E, including Glu at position 2. A polar position 2 Gln yielded the strongest binding signals for both the HLA-B7 and HLA-A1 leader peptides demonstrating a degree of consistency in hierarchies of relative binding affinity between VL9 peptide variants. Mtb44 peptide variants with a range of non-canonical primary anchor residues at positions 2 and 9 were also screened (Figure 7, B). The inclusion of charged anchor residues such as Glu or Lys at positions 2 or 9 greatly reduced binding of Mtb44 peptide variants to HLA-E, although Lys at position 2 did yield significant HLA-E binding. In addition to wild-type Mtb44, three variants demonstrated HLA-E binding with the greatest degree of statistical significance in the peptide-exchange assay, including those with position 2 or 9 Phe and position 2 Gln substitutions.

**Figure 7.**
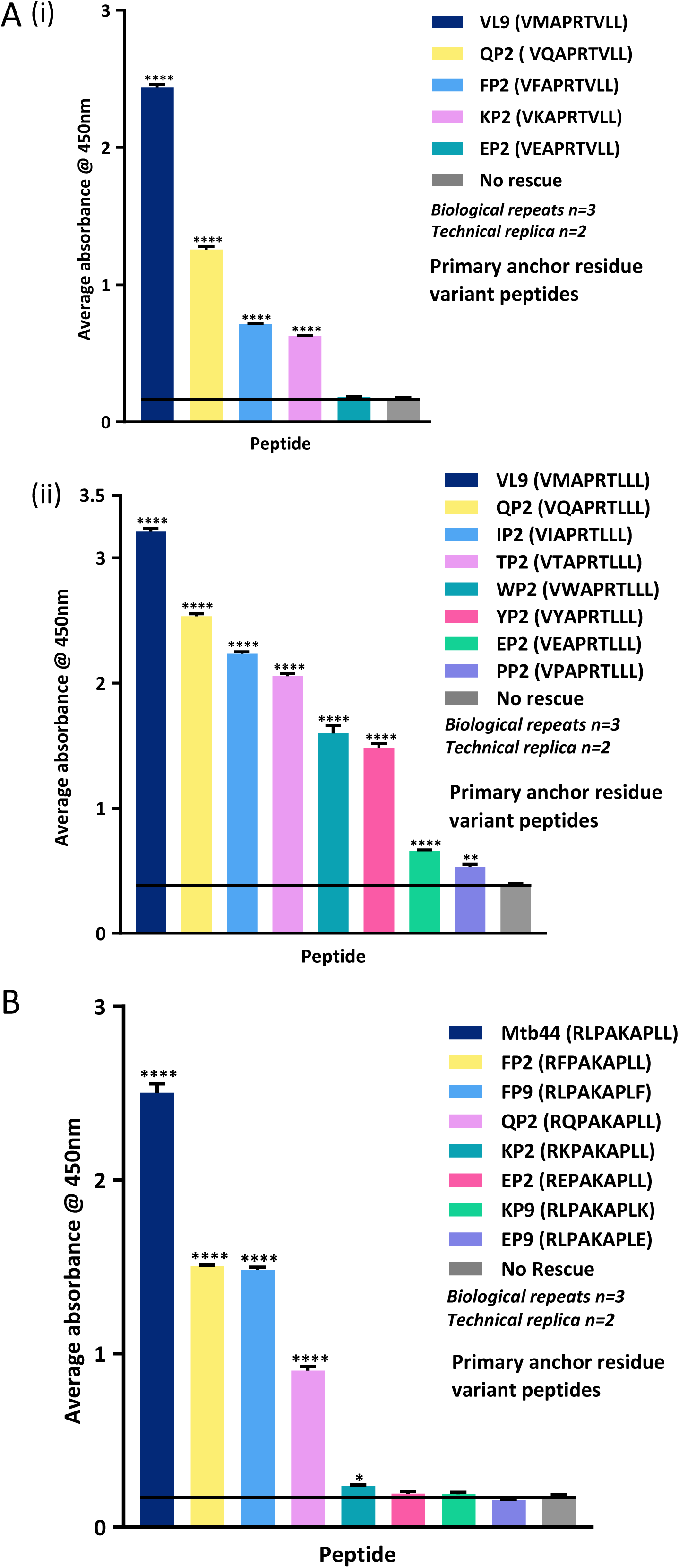
Peptide exchange ELISA binding assay screening of VL9 and Mtb44 position 2 and 9 primary anchor residue variants. **A. (i)** Average absorbance readings from the peptide-exchange sandwich-ELISA-based binding assay for a selection of VL9 position 2 variant peptides. The positive control VL9 peptide (VMAPRTVLL) is shown in dark blue with the peptide-free background (No rescue) in grey. Peptide ID, sequence, length and corresponding references are detailed in Supplementary Table 1, D (i). **(ii)** Average absorbance readings from the peptide-exchange sandwich-ELISA-based binding assay are denoted on the Y-axis for a selection of VL9 position 2 variant peptides based on the sequence, VMAPRTLLL. Peptide details are listed in Supplementary Table 1, D (i). **B.** Average absorbance readings from the peptide-exchange sandwich-ELISA-based binding assay for a collection of Mtb44 primary anchor variant peptides with a range of alternative hydrophobic, polar or charged residues in place of the canonical primary anchor residues. The positive control wild-type Mtb44 peptide (RLPAKAPLL) is shown in dark blue with the peptide-free background (No rescue) in grey. Average absorbance readings at 450nm are displayed on the Y-axis for plots in **A** and **B**. Peptide details and corresponding references are denoted in Supplementary Table 1, D (ii). Error bars depicting the mean and standard error of the mean (SEM) of biological repeats are shown.

### ELISA-based HLA-E peptide binding sequence motif

ELISA signals for peptides screened in the peptide exchange assay were normalised via background (no-peptide rescue) subtraction and subsequently expressed as percentages of the positive control VL9 peptide signal enabling the establishment of an inter-assay hierarchy of peptide binding to HLA-E presented as a heat map in Figure 8 A (i). Peptides were ranked in order of relative binding strength to HLA-E according to ELISA data and subdivided into 11 categories from those that generated signals below background levels to those that generated signals ranging from 70-100% of the positive control signal following background subtraction (Supplementary Table 2). A Seq2Logo HLA-E sequence motif ^31^ was generated based on nonameric peptide sequences which generated signals 20-100% of VL9 following buffer subtraction (Figure 8, A, ii). VL9 (VMAPRTVLL/VMAPRTLLL) and Mtb44 (RLPAKAPLL) primary anchor variant peptides were excluded from this analysis to avoid overrepresentation of residues at remaining peptide positions. Unsurprisingly, primary anchor positions 2 and 9 imposed the greatest degree of specificity, as indicated by their over-representation relative to the remaining residue positions in the Seq2Logo motif. Further, the most frequently occurring residue at position 9, is consistent with canonical position 9 Leu. Leu was also the most frequently observed residue at position 2 of the Seq2Logo followed by canonical Met. A range of other hydrophobic residues were also tolerated at both primary anchor positions including Phe, Val, Ala and Ile, in addition to a polar Gln at position 2. Interestingly, Pro was the most frequently occurring residue at positions 3, 4, 6 and 7 of the Seq2Logo motif perhaps indicating a preference for this torsion angle-restricted residue at a central position along HLA-E-binding peptides, especially when preferred secondary anchor residues are absent. Additionally, residues with small side chains such as Ala and Gly were also enriched at central peptide positions 3, 4, 6 and 7. Unsurprisingly, due to its solvent exposed orientation in VL9-bound structures of HLA-E, position 5 exhibited the shortest stack length in the Seq2Logo reflecting a greater tolerance for biochemically diverse residues at this position. In summary, it appears that HLA-E-restricted peptides can contain a range of hydrophobic residues at both primary anchor positions and are often enriched for a central Pro at positions 3, 4, 6 or 7.

**Figure 8.**
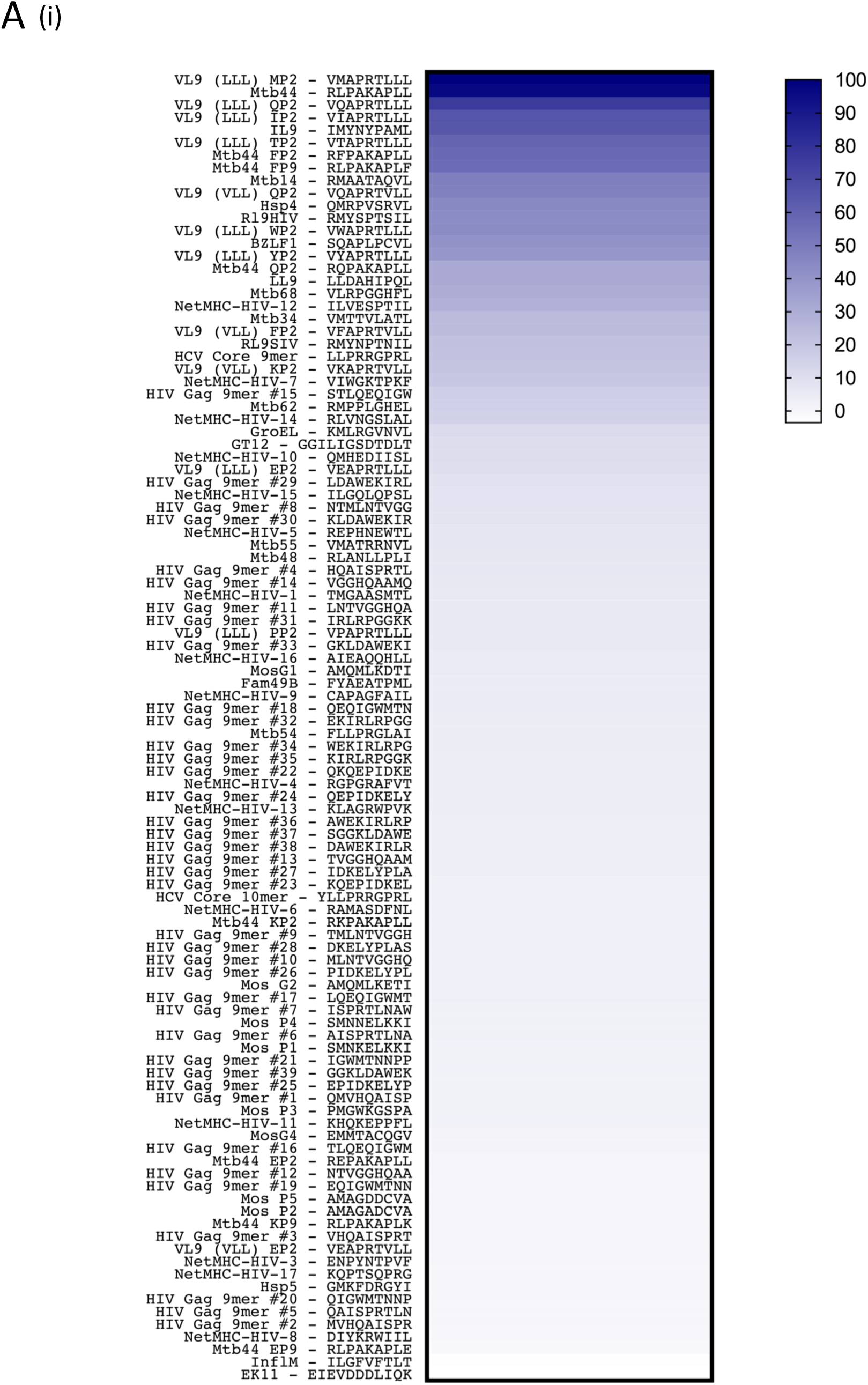
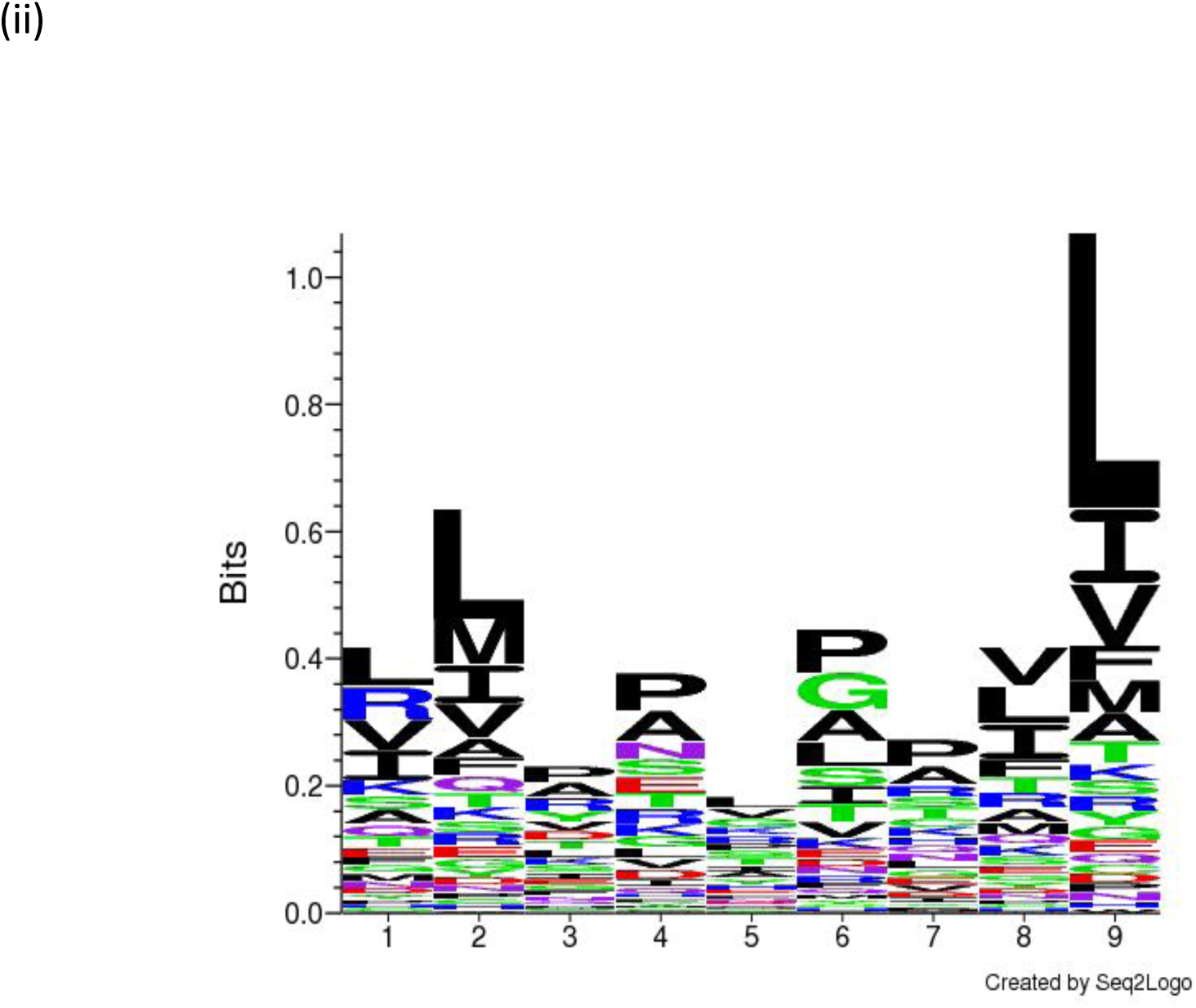
Hierarchy of peptide binding to HLA-E and Seq2Logo peptide binding motif. **A. (i)** A heat map depicting peptides screened in the peptide exchange ELISA-assay in descending order of relative binding strength to HLA-E. Screened peptides were ranked in order of binding strength to HLA-E by expressing each peptide signal as a percentage of the positive control VL9 signal following background subtraction to normalise for inter-assay variation. Each peptide is listed in Supplementary Table 1 along with its sequence, ID and corresponding percentage of VL9 signal after background subtraction. **(ii)** A Seq2Logo HLA-E peptide sequence motif based on statistically significant nonameric peptides that generated peptide exchange ELISA-based assay signals within 20-100% of the positive control VL9 signal after background subtraction. Primary anchor residue variant peptides based on either VL9 or Mtb44 peptide backgrounds were excluded from analysis to remove bias from overrepresented residues at remaining non-mutated positions. The Y axis denotes information in bits with peptide position plotted on the X-axis. Letters represent amino acids with font size indicating residue frequency. The length of residue stacks indicates the degree of conservation among inputted sequences at that position with small letter stack lengths reflecting increased residue variability.

## Discussion

Recent evidence demonstrating that HLA-E and the rhesus homologue, Mamu-E, present sequence diverse, non-VL9-like pathogen-derived peptides to CD8^+^ T cells in *Mycobacteria tuberculosis* (Mtb) infection and Cytomegalovirus-vectored SIV vaccination, respectively, suggests an unanticipated role for HLA-E in the context of T cell immunity. This previously undiscovered role is exciting and offers potential for ‘universal’ HIV vaccine development. Central to this is a requirement to develop sensitive tools to re-appraise the peptide binding ligandome of HLA-E. Consequently, we have designed and optimised a highly sensitive peptide exchange ELISA-based assay enabling the relative quantification of peptide binding to HLA-E. Comparative assays demonstrated this new peptide exchange technique offers superior sensitivity and reliability relative to the previously published micro-refolding HLA-E peptide binding ELISA ^7 20^.

Regarding assay optimisation, the identification of peptide exchange independent of photo-illumination by the dedicated UV cabinet raises the question of whether the photo-labile VL9-based peptide (VMAPJTLVL) is cleaved by lower wavelengths in the visible spectrum or by background UV radiation present in the lab. Alternatively, it is possible that full length, nonameric UV-labile peptide dissociates from the HLA-E binding groove over time in the absence of rescue peptide, irrespective of exposure to electromagnetic radiation. We previously showed, via blue native gel analysis of HLA-E, that a molar excess of lower affinity peptide (RL9HIV) can facilitate peptide exchange with a pre-bound higher affinity VL9 peptide ^20^. This demonstrates that peptide exchange from a higher to lower affinity peptide ligand is possible for HLA-E in the absence of a photo-cleavable peptide or UV irradiation when the lower affinity peptide is in molar excess. Further, the introduction of an unnatural beta amino acid at position 5 of the UV-labile VL9 peptide results in conformational changes to the peptide main chain which can in turn impact binding affinity ^23^. Therefore the modified VL9 peptide might have an even greater propensity for dissociation and consequent peptide exchange relative to wild-type VL9.

Despite various optimisation strategies improving both reliability and sensitivity of the peptide exchange ELISA assay, some technical challenges remain. First, inter-assay variation may be introduced due to stability differences in macro-refolded HLA-E-β2M-VMAPJTVLL complexes. Since the UV-labile VL9 ligand appears to dissociate from pre-refolded HLA-E complexes in the absence of UV-illumination, the precise timings of size exclusion chromatography, purified protein concentration and subsequent freezing methods could influence the degree of peptide dissociation prior to the peptide exchange reaction, and subsequently impact ELISA data. To account for such inter-assay variation, we normalised average absorbance readings of test peptides by subtracting the background signal and subsequently expressing binding signals as percentages of VL9 binding before generating an overall inter-assay hierarchy of peptide binding to HLA-E. Additionally, VL9 positive control peptides generate much higher average absorbance readings than the majority of screened peptides and these positive control signals remain unaffected when the VL9 peptide concentration is titrated from 100 to below 10µM (Supplementary Figure 2). This, coupled with increased assay sensitivity, renders sample spacing on ELISA plates a crucial factor to consider to avoid cross-contamination. We observed that ELISA wells containing the VL9 positive control must be separated by at least two wells in each direction or ideally situated on a separate plate to avoid this issue (data not shown). This also extends to medium binding peptides that generate signals over 10% of the VL9 positive control following background subtraction – such peptides must also be separated by multiple wells from lower affinity, non-binding peptides or negative controls to eliminate false positive results.

The optimised peptide exchange ELISA assay reliably demonstrated HLA-E binding to a range of pathogen- and self-derived peptides with sequences that deviate from the canonical VL9 peptide. Such results support previous findings that in certain physiological contexts the HLA-E peptide binding repertoire might not be limited to MHC class I-derived leader peptides ^17^. Despite this, our results are consistent with the original characterisation of the HLA-E peptidome, in which a strong preference for VL9 peptides was established ^6 1^. Of all the screened peptides, only one (Mtb44, RLPAKAPLL) exhibited comparable binding signals to the positive VL9 peptide - the Mtb44 signal represented 97% of VL9 signal following background subtraction (Supplementary Table 2). The majority of peptides tested exhibited exceptionally low binding signals many of which were indistinguishable from the negative control peptide-free background. However, a small selection of pathogen- or self-derived non-VL9 peptides demonstrated reliable, statistically significant binding to HLA-E and generated average absorbance readings above 10% of the VL9 control following background subtraction (Supplementary Table 2). The majority of these >10% of VL9 ‘medium’ strength HLA-E-binders contained the canonical primary anchor residues at positions 2 (Met) and 9 (Leu). However, BZLF1 (SQAPLPCVL) contains a polar Gln at position 2 in place of the hydrophobic Met and the peptide termed NetMHC-HIV-7 (VIWGKTPKF) contains a larger aromatic Phe at position 9 and an Ile at position 2. Our previous blue native gel analysis established that peptides, such as RL9HIV (RMYSPTSIL), within the medium affinity category which reliably generate the greatest peptide to background ratios other than VL9 or Mtb44 (RLPAKAPLL) in the ELISA assay, produce multiple gel bands on blue native gel analysis, indicative of heterogeneous protein populations ^20^. Thus, HLA-E-restricted peptides in the medium affinity category established in this study are unlikely to support formation of homogenous fully-folded HLA-E forms. However, we were previously able to obtain crystal structures of HLA-E bound to medium affinity peptides, including RL9HIV (RMYSPTSIL), indicating that a subpopulation of fully folded material was incorporated into the crystal lattice ^20^. Previously obtained peptide-bound structures of HLA-E and their corresponding ELISA rankings are listed in Supplementary Table 3. Although such structures were obtained for HLA-E binding peptides that generated ELISA signals between 31.3% and 96.5% of the VL9 positive control following buffer subtraction, crystals could not be grown for lower binding peptides likely owing to increased complex instability inhibiting lattice formation. Crystals also could not be obtained for the BZLF EBV-derived peptide (SQAPLPCVL) despite its ELISA ranking (40.7% of VL9) falling within range of other crystallisable peptides.

To further probe the tolerability of the primary anchor B and F pockets of the HLA-E binding groove we synthesised VL9- and Mtb44-based peptide variants and screened these primary anchor mutant peptides in the peptide exchange ELISA assay (Figure 7). A number of Mtb44 and VL9 variant peptides supported HLA-E binding despite the presence of non-canonical primary anchor residues such as a Thr, Ile, Gln, Tyr, Trp, Phe or positively charged Lys at position 2 of VL9. Notably, the hierarchy of relative binding of position 2 anchor residue mutations differed on the Mtb44 and VL9 peptide panel backgrounds, indicating other positions along these nonamers operate synergistically with position 2 substitutions to determine overall peptide binding affinity. For example, position 2 Lys generated low ELISA signals when substituted into the Mtb44 peptide backbone but yielded stronger binding when inserted into the VL9 nonamer. Furthermore, Gln at position 2 of the VL9 peptide yielded stronger binding to HLA-E compared to the position 2 Phe VL9 peptide variant. However, this trend was reversed for Mtb44 with the position 2 Phe variant demonstrating stronger binding to HLA-E, above that of the position 2 Gln variant. The presence of non-canonical anchor residues in HLA-E binding peptides does not however necessarily indicate broadly receptive pockets in the groove. Structural evidence would be required to elucidate whether non-canonical primary anchor side chains occupy their corresponding pockets, or whether these peptides adopt alternative binding conformations.

Early experiments delineating the HLA-E binding motif analysed acid eluted peptide from *in vitro* refolded HLA-E with VL9 anchor residue variant peptide libraries via mass spectrometry ^32^. However, the HLA-E binding groove’s capacity to accommodate suboptimal peptides and the sequence identity of potential non-VL9 HLA-E-binding peptides was not fully elucidated. For example, the sensitivity of peptide elution from *in vitro* refolded HLA-E would have been compromised by the co-presence of an excess of near optimal VL9 variant peptide in the pool - such as those with position 2 Met to Leu or Ile substitutions which were shown to only marginally reduce binding affinity to HLA-E ^32^. Peptides with lower binding affinities would likely have been outcompeted in these settings and thus remain undetected. By contrast, the peptide exchange ELISA assay presented here affords superior sensitivity as the binding capacity of suboptimal HLA-E binding peptides is tested in the absence of competition from higher affinity peptide. Although previous mass spectrometry studies identified an extended hydrophobic primary anchor motif for HLA-E that included residues such as Ile and Leu in addition to Met at position 2, our motif analysis builds on previous work. The HLA-E peptide binding motif based on screens presented here, demonstrates that additional hydrophobic residues such as Val, Ala and Phe can be tolerated at both primary anchor positions in addition to a polar Gln at position 2 (Figure 8, A, ii). This extended motif is supported by our previously published structures of HLA-E in complex with Mtb44 (RLPKAPLL) primary anchor variant peptides containing a Phe at position 2, Phe at position 9 and Gln at position 2 ^20^. All substituted primary anchor side chains were accommodated by their corresponding primary B and F pockets and superposed to the wild-type anchor side chains demonstrating increased primary pocket tolerability. Beyond this, our ELISA-based HLA-E sequence motif also highlighted the importance of a central Pro at positions 3, 4, 6 or 7. Interestingly, peptides which satisfied the primary anchor motif demonstrated a large range of binding strength for HLA-E in the peptide exchange ELISA assay thus illustrating that the presence of particular hydrophobic residues at positions 2 and 9 is not sufficient for optimal HLA-E binding. However, ten of the twelve non-VL9 pathogen- or self-derived peptides (excluding Mtb44 and VL9 primary anchor variant peptides) that generated signals >20% of VL9 following background subtraction contained a Pro at a central position in the peptide, in addition to tolerated primary anchors. For example, Mtb44 (RLPAKAPLL), the strongest binding peptide tested, contains a Pro at both positions 3 and 7. Two of these medium binding peptides (>20% of VL9), RL9HIV and RL9SIV, contain a Pro at the largely unrestricted position 5 as opposed to position 3, 4, 6 or 6, as indicated by the novel HLA-E sequence motif. However, the alternative peptide backbone conformation observed in the structure of RL9HIV-bound HLA-E causes the position 5 Pro side chain to align to position 6 of VL9 peptides, project towards the groove floor and thereby act as a compensatory position 6 secondary anchor residue ^33^. In this way, the Pro at position 5 of RL9HIV and RL9SIV peptides is equivalent to position 6 in other HLA-E binding peptides which adopt the classical kinked peptide configuration and contain a largely solvent exposed position 5 which projects away from the binding groove (such as Mtb44, RLPAKAPLL) ^33^. Thus, the RL9 peptides functionally adhere to the Seq2Logo motif. By contrast, the vast majority of test peptides which satisfy the primary anchor motif and generated signals <5% of VL9 following background subtraction, including NetMHC-HIV-8 (DIYKRWIIL), Hsp5 (GMKFDRGYI), HIV Gag 9mer #16 (TLQEQIGWM), MosG4 (EMMTACQGV), MosP1 (SMNKELKKI), MosP4 (SMNNELKKI), MosG2 (AMQMLKETI), HIV Gag 9mer #26 (PIDKELYPL), NetMHC-HIV-6 (RAMASDFNL), HIV Gag 9mer #13 (TVGGHQAAM), MosG1 (AMQMLKDTI) and NetMHC-HIV-16 (AIEAQQHLL), lacked a centrally-positioned Pro illustrating the importance of this additional component of the optimal HLA-E peptide sequence motif, particularly in the absence of preferred secondary anchor residues.

A Met to Thr substitution present at position 2 in a subset of HLA-B27 (VTAPRTLLL) and HLA-B15 (VTAPRTVLL) leader peptides has previously been shown to reduce HLA-E cell surface expression due to decreased peptide affinity and complex stability ^2 4^. Thus, it could be predicted that other pathogen- or self-derived peptides with lower peptide exchange ELISA assay rankings to that of VL9 position 2 Thr (Supplementary Table 2), might also exhibit compromised cell surface expression and reduced representation in the steady-state HLA-E-presented repertoire. However, the primary contexts in which HLA-E- or Mamu-E-restricted epitopes have been identified involve deregulated MHC class I trafficking pathways ^17 21 18^. It is therefore possible that perturbations in the supply of MHC class I-derived VL9 peptides in addition to alternative trafficking due to Mtb- or CMV vector-driven immune evasion mechanisms, give rise to the emergence of lower affinity, non-canonical peptides in the cell surface-presented HLA-E ligandome.

Intriguingly a number of key epitopes against which HLA-E-restricted CD8^+^ T cell responses have previously been identified did not exhibit definitive HLA-E binding in this assay. For example, the Mtb-derived EK11 which comprises residues 19-29 of the conserved hypothetical protein Rv0634A (EIEVDDDLIQK), demonstrated an immunodominant profile with HLA-E-restricted responses detected in 81% of tested donors with active or latent Mtb infection ^21^. However, in the peptide exchange ELISA assay EK11 generated an average absorbance reading below the ‘no-rescue’ peptide-free background signal, and produced the lowest signal of any peptide tested (Supplementary Table 2). Further, a Fam49B-derived peptide (FYAEATPML) previously shown to elicit Qa-1-restricted CD8^+^ T cell responses also demonstrated exceptionally low signals below 5% of the VL9 positive control signal following buffer subtraction in our assay system ^14^. Such results are consistent with previous micro-refolding ELISA and single chain trimer transduction data which indicate the 11 optimal SIV epitopes in RhCMV 68-1 vector-based vaccine studies also bind HLA-E with dramatically lower relative affinity than the positive control VL9 peptide, with some generating signals only marginally above background ^17^. Surprisingly, these epitopes appear to play an important and protective physiological role despite not supporting HLA-E complex stability in this assay system. It therefore appears that peptide binding affinity for HLA-E may not always correlate with the strength of the CD8^+^ T cell response elicited and that perhaps two modes of HLA-E-restricted CD8^+^ T cell recognition exist; the first of which comprises conventional recognition of medium strength HLA-E-restricted peptides and the second of which tentatively involves exceptionally low affinity or apparent non-binders. Alternatively, the conditions of our *in vitro* assay may not adequately reflect the intracellular environments that support MHC-E peptide loading in RhCMV 68-1 vaccinated or Mtb-infected cells. Equally, it is possible that the conventional mode of peptide binding might not apply in these settings but instead a currently unknown mechanism could facilitate suboptimal peptide presentation or stabilise low affinity peptide binding to MHC-E. Abacavir, an anti-retroviral nucleoside analogue reverse transcriptase inhibitor used to treat HIV-infected individuals, was structurally demonstrated to specifically bind to the C/F pockets of HLA-B*5701, in turn altering the presented peptide repertoire and resulting in self-reactivity ^34^. It is possible that, analogous to this drug-modified repertoire, pocket stabilising interactions with an unidentified co-factor might exclusively occur in contexts such as RhCMV 68-1-SIV vaccination or Mtb infection, thus sufficiently stabilising HLA-E complexes to permit the presentation of low affinity peptide or alter the landscape of the HLA-E binding groove to transform its peptide binding preferences.

In summary, the highly sensitive peptide exchange ELISA-based binding assay presented here can reliably establish the relative HLA-E binding capacity of test peptides. Considering HLA-E-, Qa-1- and Mamu-E-restricted CD8^+^ T cell responses have been identified in various contexts of vaccination, infection and autoimmune disease, this assay provides a useful tool to validate and determine binding strength of putative HLA-E-restricted epitopes. Further, this assay can be used to screen algorithmically-identified peptides predicted to support HLA-E complex stability, thus potentially enabling the identification of novel HLA-E-restricted peptides which could subsequently be targeted in therapeutic or vaccine settings. In future, it will be important to determine the underlying mechanistic basis of MHC-E-restricted CD8^+^ T cell recognition and resolve how such remarkably low affinity or apparent ‘non-binding’ peptides elicit robust HLA-E-, Qa-1- or Mamu-E-restricted CD8^+^ T cell responses *in vivo* ^17^.

## Materials and Methods

### Peptide synthesis

Synthetic peptides were generated by Fmoc (9-fluorenylmethoxy carbonyl) chemistry to a purity of 85% by Genscript USA. All peptides were provided as lyophilised powder, reconstituted in DMSO to a concentration of 200mM, and stored at -80 °C. A UV photo-labile version of the HLA-B leader sequence peptide, VMAPRTLVL, incorporating a UV-sensitive 3-amino-3-(2-nitrophenyl)-propionic acid residue (J residue) substitution at position 5, VMAPJTLVL was synthesised by Dris Elatmioui at LUMC, The Netherlands. The UV-sensitive peptide (termed 7MT2) was stored as lyophilised powder, and reconstituted as required.

### HLA-E protein refolding and purification

β2-microglobulin in Urea-Mes, at a final concentration of 2μM, was refolded at 4 °C for 30mins in a macro-refolding buffer prepared in MiliQ water containing 100 mM Tris pH8.0, 400mM L-arginine monohydrochloride, 2mM EDTA, 5mM reduced glutathione and 0.5mM oxidised Glutathione. The UV-sensitive peptide 7MT2 was subsequently added to the β2M refold to achieve a peptide concentration of 30μM. HLA-E*0103 heavy chain was subsequently pulsed into the refolding buffer until a final concentration of 1μM was reached. Following incubation for 72hrs at 4 °C, HLA-E refolds were filtered through 1.0μm cellular nitrate membranes to remove aggregates prior to concentration by the VivaFlow 50R system with a 10kDa molecular weight cut-off (Sartorius) and subsequent concentration in 10kDa cut-off VivaSpin Turbo Ultrafiltration centrifugal devices (Sartorius). Samples were then separated according to size into 20 mM Tris pH8, 100 mM NaCl by fast protein liquid chromatography (FPLC) on an AKTA Start System using a Superdex S75 16/60 column. Elution profiles were visualised by UV absorbance at 280mAU, enabling differentiation of correctly refolded HLA-E-β2M-pepide complexes from smaller non-associated β2M and larger misfolded aggregates. FPLC-eluted protein peak fractions were combined and concentrated to 3 mg/mL using 10kDa cut-off VivaSpin Turbo Ultrafiltration centrifugal devices prior to use in the HLA-E peptide exchange ELISA-based assay. 1μL aliquots were analysed by non-reducing SDS-PAGE electrophoresis on NuPAGE™ 12% Bis-Tris protein gels (ThermoFisher Scientific) to confirm the presence of non-aggregated HLA-E heavy chain and β2m.

### HLA-E peptide exchange reaction

3µg of purified HLA-E previously refolded with the UV sensitive peptide 7MT2 was incubated in the presence of 100 µM ‘exchange’ peptide in polypropylene V-shaped 96-well plates (Greiner Bio-One) for 5 hours on ice. The final volume of each well was adjusted to 125μL by adding exchange buffer comprising 100 mM Tris pH8.0, 400mM L-arginine monohydrochloride, 2mM EDTA, 5mM reduced glutathione and 0.5mM oxidised Glutathione, prepared in MiliQ water. The optimisation of certain variables of this technique including buffer composition, duration of incubation and UV exposure is detailed in the text of the results section where the optimisation strategy of the HLA-E peptide exchange ELISA-based assay is discussed.

### Sandwich ELISA

96 well ELISA plates (Life Technologies) were coated with the anti-human HLA-E monoclonal capture antibody, 3D12 (BioLegend), diluted to 10μg/mL in freshly prepared ELISA Coating Buffer (BioLegend) and incubated for 12 hours at 4°C. Wells were washed three times in 200μL phosphate buffered saline (PBS), and subsequently incubated in 300μL of 2% IgG-free bovine serum albumin (BSA) (Sigma Aldrich) for 2 hours at room temperature to block unoccupied surfaces on the ELISA plate wells to prevent non-specific protein binding. Following plate blocking, plates were washed five times with 200µL freshly prepared 0.05% Tween-based ELISA Wash Buffer (BioLegend) and a final time in 200μL phosphate buffered saline (PBS) (all subsequent plate washing was carried out in an identical manner). 50μL of the HLA-E reaction mixture from either micro-scale refolding or UV-induced peptide exchange, diluted 1:100 in 2% BSA, was added to each well of the ELISA plate and left to incubate for 1 hour at room temperature. Following protein incubation and subsequent plate washing, 50μL of polyclonal anti-human β2M HRP-conjugated IgG detection antibody raised in rabbits (ThermoFisher Scientific), diluted 1:2500 in 2% BSA (0.2μg/mL), was added to each well to ensure detection of β2M-associated forms of HLA-E only. After 30mins of dark incubation at 4°C, ELISA plates were washed and 50µL of a goat anti-rabbit HRP-conjugated IgG enhancement antibody (EnVision+ System-HRP from Agilent), diluted 1:15 in 2% BSA containing 1% mouse serum to prevent cross-reactivity, was added to each well to amplify the signal. ELISA plates were incubated for 15mins in the dark at room temperature, and following a final round of plate washing, developed in 100μL of 3,3’,5,5’-tetramethyl benzidine (TMB) substrate (BioLegend) and incubated in the dark at room temperature for a further 10 minutes. Finally, the reactions were terminated in 100μL of H_2_SO_4_ STOP solution (BioLegend) and absorbance readings were obtained at 450nm on a FLUOstar OMEGA plate reader. Both the peptide exchange reaction and sandwich ELISA stages of the assay are depicted in Supplementary Figure 1.

## Supporting information

All Suppl Figures and Files

## Acknowledgements

This work was supported by grants from the BMGF OPP1133649, MRC MR/M019837/1, NIAID 5UM1AI126619-05 and the Hester Cordelia Parsons Fund, Oxford University.

