## Supplementary material for "Detailed and atypical HLA-E peptide binding motifs revealed by a novel peptide exchange binding assay": All Suppl Figures and Files

### Supplementary Information

A novel peptide exchange binding assay reveals both detailed and atypical motifs for peptides that bind HLA-E

***Lucy C. Walters et al.***

#### Supplementary Figure 1

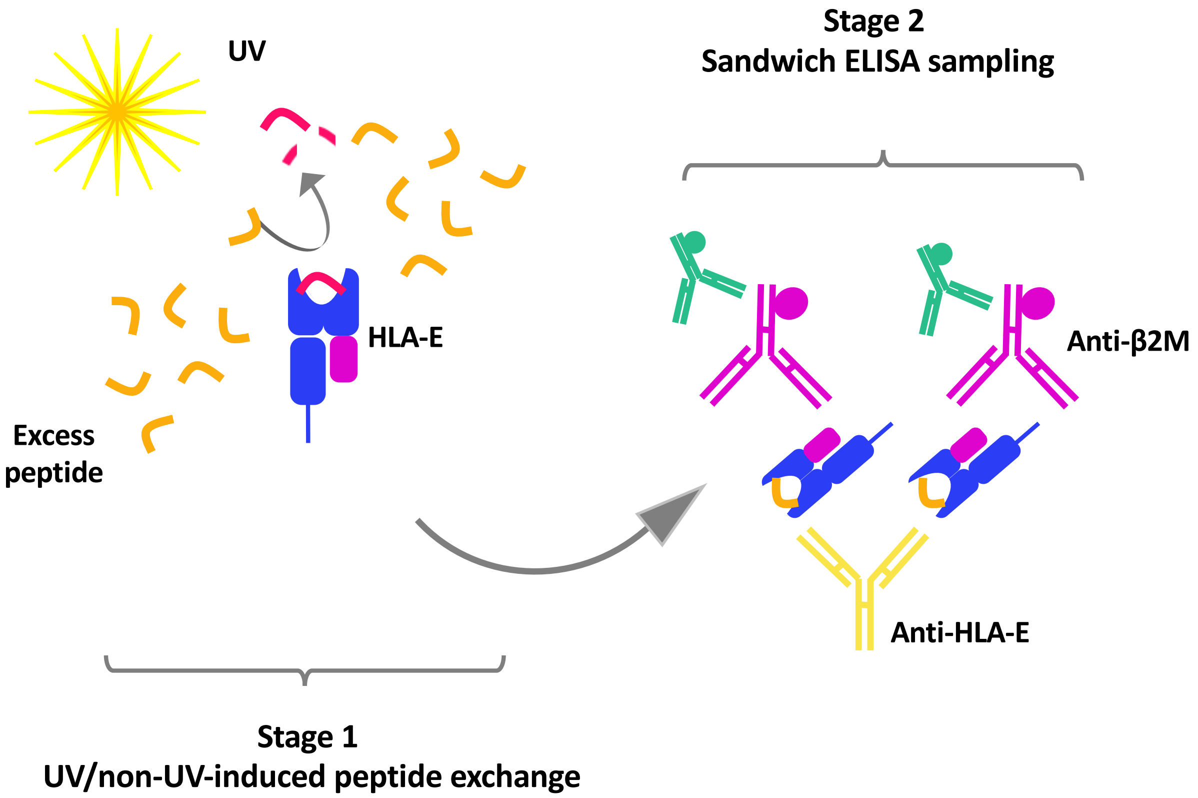


**Supplementary Figure 1. UV/non-UV-induced peptide exchange ELISA-based method visualisation**
In stage 1 of the UV/non-UV-induced peptide exchange reaction, β2M-bound (violet) HLA-E (blue) pre-refolded and purified with the UV-labile VL9 variant peptide (VMAPJTVLL) (magenta) is exposed to UV irradiation in the presence of excess test peptide. **In the presence of UV-irradiation the photo-sensitive VL9 peptide is cleaved and dissociates from the binding groove, also, this peptide dissociates independent of photo-induced cleavage**. A molar excess of test peptide (orange) enables peptide exchange. The peptide binding affinity of the rescue peptide (orange) determines the concentration of correctly refolded β2M-bound HLA-E complexes in solution and in the absence of a rescue peptide the complexes dissociate. The relative concentration of β2M-bound HLA-E complexes is determined indirectly via sandwich ELISA whereby a sample of the peptide exchange reaction is added to an ELISA plate pre-coated with the HLA-E-specific antibody 3D12 (yellow) in stage 2 of the assay. The subsequent addition of a horse radish peroxidase-coupled β2M-specific detection antibody (violet) and subsequent antibody-enabled amplification (green) enables the detection of β2M-bound HLA-E complexes-only. Reactions are developed with TMB and terminated with STOP solution prior to obtaining absorbance signals at 450nm.

### Supplementary Table 1

## A (i)

Peptides predicted to bind HLA-E by NetMHC

| Peptide ID | Sequence | Aas | Organism | Protein | Ref |
|---|---|---|---|---|---|
| NetMHC-HIV-1 | TMGAASMTL | | | Env | |
| NetMHC-HIV-2 | HQPISARTL | | | Gag/pol | |
| NetMHC-HIV-3 | ENPYNTPVF | | | Gag/pol | |
| NetMHC-HIV-4 | RGPGRAFVT | | | Env | |
| NetMHC-HIV-5 | REPHNEWTL | | | Vpr | |
| NetMHC-HIV-6 | RAMASDFNL | | | Gag/pol | |
| NetMHC-HIV-7 | VIWGKTPKF | | | Gag/pol | |
| NetMHC-HIV-8 | DIYKRWIIL | | | Gag/pol | |
| NetMHC-HIV-9 | CAPAGFAIL | 9 | HIV | Env | Andreatta M & Nielsen M. Bioinformatics (2016) |
| NetMHC-HIV-10 | QMHEDIISL | | | Env | |
| NetMHC-HIV-11 | KHQKEPPFL | | | Gag/pol | |
| NetMHC-HIV-12 | ILVESPTIL | | | Rev | |
| NetMHC-HIV-13 | KLAGRWPVK | | | Gag/pol | |
| NetMHC-HIV-14 | RLVNGSLAL | | | Env | |
| NetMHC-HIV-15 | ILGQLQPSL | | | Gag/pol | |
| NetMHC-HIV-16 | AIEAQQHLL | | | Env | |
| NetMHC-HIV-17 | KQPTSQPRG | | | Tat | |

## (ii)

HIV-derived peptides with optimal position 2 Met

| Peptide ID | Sequence | Aas | Organism | Protein | Ref |
|---|---|---|---|---|---|
| MosG1 | AMQMLKDTI | | | | |
| MosG2 | AMQMLKETI | | | Gag | |
| MosG4 | EMMTACQGV | | | | |
| MosP1 | SMNKELKKI | | | | Bette Korber, unpublished |
| MosP2 | AMAGADCVA | 9 | HIV | | |
| MosP3 | PMGWKGSPA | | | Pol | |
| MosP4 | SMNNELKKI | | | | |
| MosP5 | AMAGDDCVA | | | | |

(iii)

**Overlapping HIV Gag 9mers**

| Peptide ID | Sequence | Aas | Organism | Protein |
|---|---|---|---|---|
| HIV Gag 9mer #1 | QMVHQAISP | | | |
| HIV Gag 9mer #2 | MVHQAISPR | | | |
| HIV Gag 9mer #3 | VHQAISPRT | | | |
| HIV Gag 9mer #4 | HQAISPRTL | | | |
| HIV Gag 9mer #5 | QAISPRTLN | | | |
| HIV Gag 9mer #6 | AISPRTLNA | | | |
| HIV Gag 9mer #7 | ISPRTLNAW | | | |
| HIV Gag 9mer #8 | NTMLNTVGG | | | |
| HIV Gag 9mer #9 | TMLNTVGGH | | | |
| HIV Gag 9mer #10 | MLNTVGGHQ | | | |
| HIV Gag 9mer #11 | LNTVGGHQA | | | |
| HIV Gag 9mer #12 | NTVGGHQAA | | | |
| HIV Gag 9mer #13 | TVGGHQAAM | | | |
| HIV Gag 9mer #14 | VGGHQAAMQ | | | |
| HIV Gag 9mer #15 | STLQEQIGW | | | |
| HIV Gag 9mer #16 | TLQEQIGWM | | | |
| HIV Gag 9mer #17 | LQEQIGWMT | | | |
| HIV Gag 9mer #18 | QEQIGWMTN | | | |
| HIV Gag 9mer #19 | EQIGWMTNN | | | |
| HIV Gag 9mer #20 | QIGWMTNNP | 9 | HIV | Gag |
| HIV Gag 9mer #21 | IGWMTNNPP | | | |
| HIV Gag 9mer #22 | QKQEPIDKE | | | |
| HIV Gag 9mer #23 | KQEPIDKEL | | | |
| HIV Gag 9mer #24 | QEPIDKELY | | | |
| HIV Gag 9mer #25 | EPIDKELYP | | | |
| HIV Gag 9mer #26 | PIDKELYPL | | | |
| HIV Gag 9mer #27 | IDKELYPLA | | | |
| HIV Gag 9mer #28 | DKELYPLAS | | | |
| HIV Gag 9mer #29 | LDAWEKIRL | | | |
| HIV Gag 9mer #30 | KLDAWEKIR | | | |
| HIV Gag 9mer #31 | IRLRPGGKK | | | |
| HIV Gag 9mer #32 | EKIRLRPGG | | | |
| HIV Gag 9mer #33 | GKLDAWEKI | | | |
| HIV Gag 9mer #34 | WEKIRLRPG | | | |
| HIV Gag 9mer #35 | KIRLRPGGK | | | |
| HIV Gag 9mer #36 | AWEKIRLRP | | | |
| HIV Gag 9mer #37 | SGGKLDAWE | | | |
| HIV Gag 9mer #38 | DAWEKIRLR | | | |
| HIV Gag 9mer #39 | GGKLDAWEK | | | |

(iv)

| Peptide ID | Sequence | Aas | Organism | Protein | Ref. |
|---|---|---|---|---|---|
| $RL9SIV_{supertope}$ | RMYNPTNIL | 9 | *Simian immunodeficiency virus* | $Gag_{276-284}$ | Hansen, S,G. et al (2016) |
| RL9HIV | RMYSPTSIL | | *Human immunodeficiency virus* | $Gag_{275-283}$ | Walters, L,C. et al (2018) |

B (i)

Mtb-derived peptides eluted from infected cells

| Peptide ID | Sequence | Aas | Organism | Protein | Ref |
|---|---|---|---|---|---|
| IL9 | IMYNYPAML | 9 | *Mycobacteria* | $EsxH_{4-12}$ | McMurtrey, C. et al. (2017) |
| LL9 | LLDAHIPQL | 9 | | $EsxG_{3-11}$ | |
| GT12 | GGILIGSDTDLT | 12 | | $Lpql_{89-100}$ | |
| EK11 | EIEVDDDLIQK | 11 | | $Rv0634A_{19-29}$ | |

(ii) Algorithmically-identified peptides predicted to bind HLA-E

| Peptide ID | Sequence | Aas | Organism | Protein | Ref |
|---|---|---|---|---|---|
| Mtb 14 | RMAATAQVL | 9 | *Mycobacteria* | Rv2932 | Joosten, S.A, et al. (2010) |
| Mtb 34 | VMTTVLATL | | | Rv1734c | |
| Mtb 44 | RLPAKAPLL | | | Rv1484 | |
| Mtb 48 | RLANLLPLI | | | Rv3189 | |
| Mtb 54 | FLLPRGLAI | | | Rv0056 | |
| Mtb 55 | VMATRRNVL | | | Rv1518 | |
| Mtb 62 | RMPPLGHEL | | | Rv2997 | |
| Mtb 68 | VLRPGGHFL | | | Rv1523 | |

C (i)

| Peptide ID | Sequence | Aas | Organism | Protein | Ref. |
|---|---|---|---|---|---|
| GroEL Salmonella | KMLRGVNVL | 9 | *Salmonella enterica* serovar Typhi | $GroEL^{15-23}$ | Salerno-Goncalves, R. et al. (2004) |
| EBV BZLF1 | SQAPLPCVL | 9 | *Epstein Barr Virus* | $BZLF1^{39-47}$ | Jørgensen, P,B. et al. (2012) |
| InflM | ILGFVFTLT | 9 | *Influenza* | $Matrix^{59-67}$ | Ulbrecht, M. et al. (1998) |
| HCV Core 10mer | YLLPRRGPRL | 10 | *Hepatitis C Virus* | HCV $core^{35-44}$ | Jiang, H. et al. (2010) |
| Hsp4 Self | QMRPVSRVL | 9 | Human | $Hsp60sp^{10-18}$ | Michaëlsson, J. et al. (2002) |
| Hsp5 self | GMKFDRGYI | 9 | Human | $Hsp60.4^{216-224}$ | Michaëlsson, J. et al. (2002) |
| Fam49B protein | FYAEATPML | 9 | Human | $Fam49b^{190-198}$ | Nagarajan, N,A. et al. (2012) |

D (i)

| Peptide ID | Sequence | Aas | Organism | Protein | Ref |
|---|---|---|---|---|---|
| VL9 (WT) | VMAPRTVLL | 9 | Human | HLA-B7/8/14/39/42/48 | O'Callaghan et al. (1998) |
| QP2 | VQAPRTVLL | | | | |
| FP2 | VFAPRTVLL | | | | |
| KP2 | VKAPRTVLL | | | | |
| EP2 | VEAPRTVLL | | | | |
| VL9 (WT) | VMAPRTLLL | 9 | Human | HLA-A1 | Strong, R,K. et al. (2003) |
| QP2 | VQAPRTLLL | | | | |
| IP2 | VIAPRTLLL | | | | |
| TP2 | VTAPRTLLL | | | | |
| WP2 | VWAPRTLLL | | | | |
| YP2 | VYAPRTLLL | | | | |
| EP2 | VEAPRTLLL | | | | |
| PP2 | VPAPRTLLL | | | | |

(ii)

| Peptide ID | Sequence | Aas | Organism | Protein | Ref |
|---|---|---|---|---|---|
| Mtb44 | RLPAKAPLL | 9 | *Mycobacteria* | Rv1484 | Joosten, S.A, et al. (2010) |
| FP2 | RFPAKAPLL | | | | |
| FP9 | RLPAKAPLF | | | | |
| QP2 | RQPAKAPLL | | | | |
| KP2 | RKPAKAPLL | | | | |
| KP9 | RLPAKAPLK | | | | |
| EP2 | REPAKAPLL | | | | |
| EP9 | RLPAKAPLE | | | | |

**Supplementary Table 1. Peptides predicted to bind HLA-E screen in peptide exchange ELISA assay**
A (i) HIV-derived peptides predicted to bind HLA-E by NetMHC. (ii) HIV peptides with optimal position 2 Met primary anchor residue and a hydrophobic position 9 primary anchor residue. (iii) Overlapping 9mer peptides derived from HIV Gag. (iv) HIV/SIV Gag epitopes identified in vaccine studies in rhesus macaques using recombinant CMV vectors. B (i) Mycobacterial peptides eluted from infected cells predicted to bind HLA-E. (ii) Mycobacterial peptides that were previously predicted to bind HLA-E. C. A table of self- or pathogen-derived peptides previously predicted to bind either HLA-E or its murine homologue, Qa-1. D. (i) A panel of primary anchor residue variant peptides based on canonical VL9 peptide backgrounds. (ii) Primary anchor residue variant peptides based on the Mtb44 peptide background. *Peptide ID, sequence, length and origin are reported along with the corresponding reference. Listed peptides were screened in the peptide exchange ELISA assay for HLA-E binding capacity.*

#### Supplementary Table 2

| Peptide ID | Organism | Sequence | %VL9 | %VL9 Category |
|---|---|---|---|---|
| Mtb44 | Mycobacteria | RLPAKAPLL | 96.5 | >70% - 100% |
| VL9 (LLL) QP2 | Self | VQAPRTLLL | 76.0 | |
| VL9 (LLL) IP2 | Self | VIAPRTLLL | 65.4 | >60% - 70% |
| IL9 | Mycobacteria | IMYNYPAML | 64.5 | |
| VL9 (LLL) TP2 | Self | VTAPRTLLL | 59.1 | >50% - 60% |
| Mtb44 FP2 | Mycobacteria | RFPAKAPLL | 57.3 | |
| Mtb44 FP9 | Mycobacteria | RLPAKAPLF | 56.2 | |
| Mtb14 | Mycobacteria | RMAATAQVL | 48.4 | >40% - 50% |
| VL9 (VLL) QP2 | Self | VQAPRTVLL | 48.0 | |
| Hsp4 | Self | QMRPVSRVL | 44.5 | |
| Rl9HIV | HIV | RMYSPTSIL | 43.4 | |
| VL9 (LLL) WP2 | Self | VWAPRTLLL | 42.9 | |
| BZLF1 | EBV | SQAPLPCVL | 40.7 | |
| VL9 (LLL) YP2 | Self | VYAPRTLLL | 38.8 | >30% - 40% |
| Mtb44 QP2 | Mycobacteria | RQPAKAPLL | 31.3 | |
| LL9 | Mycobacteria | LLDAHIPQL | 31.0 | |
| Mtb68 | Mycobacteria | VLRPGGHFL | 30.0 | >20% - 30% |
| NetMHC-HIV-12 | HIV | ILVESPTIL | 28.7 | |
| Mtb34 | Mycobacteria | VMTTVLATL | 24.4 | |
| VL9 (VLL) FP2 | Self | VFAPRTVLL | 24.1 | |
| RL9SIV | SIV | RMYNPTNIL | 21.6 | |
| HCV Core 9mer | Hepatitis C Virus | LLPRRGPRL | 21.1 | |
| VL9 (VLL) KP2 | Self | VKAPRTVLL | 20.3 | |
| NetMHC-HIV-7 | HIV | VIWGKTPKF | 19.3 | >10% - 20% |
| HIV Gag 9mer #15 | HIV | STLQEQIGW | 16.0 | |
| Mtb62 | Mycobacteria | RMPPLGHEL | 15.6 | |
| NetMHC-HIV-14 | HIV | RLVNGSLAL | 15.5 | |
| GroEL | Salmonella | KMLRGVNVL | 11.2 | |
| GT12 | Mycobacteria | GGILIGSDTDLT | 10.0 | >5% - 10% |
| NetMHC-HIV-10 | HIV | QMHEDIISL | 9.9 | |
| VL9 (LLL) EP2 | Self | VEAPRTLLL | 9.6 | |
| HIV Gag 9mer #29 | HIV | LDAWEKIRL | 8.4 | |
| NetMHC-HIV-15 | HIV | ILGQLQPSL | 7.7 | |
| HIV Gag 9mer #8 | HIV | NTMLNTVGG | 7.7 | |
| HIV Gag 9mer #30 | HIV | KLDAWEKIR | 7.0 | |
| NetMHC-HIV-5 | HIV | REPHNEWTL | 6.5 | |
| Mtb55 | Mycobacteria | VMATRRNVL | 6.3 | |
| Mtb48 | Mycobacteria | RLANLLPLI | 6.2 | |
| HIV Gag 9mer #4 | HIV | HQAISPRTL | 5.9 | |
| HIV Gag 9mer #14 | HIV | VGGHQAAMQ | 5.7 | |
| NetMHC-HIV-1 | HIV | TMGAASMTL | 5.1 | |
| HIV Gag 9mer #11 | HIV | LNTVGGHQA | 5.1 | |
| HIV Gag 9mer #31 | HIV | IRLRPGGKK | 5.1 | |
| VL9 (LLL) PP2 | Self | VPAPRTLLL | 5.1 | |

*Table continued…*

| | | | | |
|---|---|---|---|---|
| HIV Gag 9mer #33 | HIV | GKLDAWEKI | 5.0 | |
| NetMHC-HIV-16 | HIV | AIEAQQHLL | 4.9 | |
| MosG1 | HIV | AMQMLKDTI | 4.6 | |
| Fam49B | Self | FYAEATPML | 4.4 | |
| NetMHC-HIV-9 | HIV | CAPAGFAIL | 4.4 | |
| HIV Gag 9mer #18 | HIV | QEQIGWMTN | 4.2 | |
| HIV Gag 9mer #32 | HIV | EKIRLRPGG | 4.1 | |
| Mtb54 | Mycobacteria | FLLPRGLAI | 4.0 | |
| HIV Gag 9mer #34 | HIV | WEKIRLRPG | 4.0 | |
| HIV Gag 9mer #35 | HIV | KIRLRPGGK | 3.7 | |
| HIV Gag 9mer #22 | HIV | QKQEPIDKE | 3.7 | |
| NetMHC-HIV-4 | HIV | RGPGRAFVT | 3.5 | |
| HIV Gag 9mer #24 | HIV | QEPIDKELY | 3.4 | |
| NetMHC-HIV-13 | HIV | KLAGRWPVK | 3.4 | |
| HIV Gag 9mer #36 | HIV | AWEKIRLRP | 3.2 | |
| HIV Gag 9mer #37 | HIV | SGGKLDAWE | 3.2 | |
| HIV Gag 9mer #38 | HIV | DAWEKIRLR | 3.1 | |
| HIV Gag 9mer #13 | HIV | TVGGHQAAM | 3.1 | |
| HIV Gag 9mer #27 | HIV | IDKELYPLA | 3.0 | |
| HIV Gag 9mer #23 | HIV | KQEPIDKEL | 2.9 | |
| HCV Core 10mer | Hepatitis C Virus | YLLPRRGPRL | 2.9 | >1% - 5% |
| NetMHC-HIV-6 | HIV | RAMASDFNL | 2.8 | |
| Mtb44 KP2 | Mycobacteria | RKPAKAPLL | 2.7 | |
| HIV Gag 9mer #9 | HIV | TMLNTVGGH | 2.7 | |
| HIV Gag 9mer #28 | HIV | DKELYPLAS | 2.7 | |
| HIV Gag 9mer #10 | HIV | MLNTVGGHQ | 2.6 | |
| HIV Gag 9mer #26 | HIV | PIDKELYPL | 2.6 | |
| Mos G2 | HIV | AMQMLKETI | 2.6 | |
| HIV Gag 9mer #17 | HIV | LQEQIGWMT | 2.6 | |
| HIV Gag 9mer #7 | HIV | ISPRTLNAW | 2.3 | |
| Mos P4 | HIV | SMNNELKKI | 2.2 | |
| HIV Gag 9mer #6 | HIV | AISPRTLNA | 2.2 | |
| Mos P1 | HIV | SMNKELKKI | 2.1 | |
| HIV Gag 9mer #21 | HIV | IGWMTNNPP | 2.1 | |
| HIV Gag 9mer #39 | HIV | GGKLDAWEK | 2.0 | |
| HIV Gag 9mer #25 | HIV | EPIDKELYP | 2.0 | |
| HIV Gag 9mer #1 | HIV | QMVHQAISP | 1.9 | |
| Mos P3 | HIV | PMGWKGSPA | 1.9 | |
| NetMHC-HIV-11 | HIV | KHQKEPPFL | 1.7 | |
| MosG4 | HIV | EMMTACQGV | 1.3 | |
| HIV Gag 9mer #16 | HIV | TLQEQIGWM | 1.1 | |
| Mtb44 EP2 | Mycobacteria | REPAKAPLL | 0.9 | |
| HIV Gag 9mer #12 | HIV | NTVGGHQAA | 0.9 | |
| HIV Gag 9mer #19 | HIV | EQIGWMTNN | 0.9 | |
| Mos P5 | HIV | AMAGDDCVA | 0.8 | |
| Mos P2 | HIV | AMAGADCVA | 0.7 | |
| Mtb44 KP9 | Mycobacteria | RLPAKAPLK | 0.7 | |
| HIV Gag 9mer #3 | HIV | VHQAISPRT | 0.7 | |
| VL9 (VLL) EP2 | Self | VEAPRTVLL | 0.6 | >0 – 1% |
| NetMHC-HIV-3 | HIV | ENPYNTPVF | 0.5 | |
| NetMHC-HIV-17 | HIV | KQPTSQPRG | 0.5 | |
| Hsp5 | Self | GMKFDRGYI | 0.5 | |
| HIV Gag 9mer #20 | HIV | QIGWMTNNP | 0.4 | |
| HIV Gag 9mer #5 | HIV | QAISPRTLN | 0.3 | |
| HIV Gag 9mer #2 | HIV | MVHQAISPR | 0.2 | |
| NetMHC-HIV-8 | HIV | DIYKRWIIL | -0.3 | |
| Mtb44 EP9 | Mycobacteria | RLPAKAPLE | -0.9 | |
| InflM | Influenza | ILGFVFTLT | -3.5 | Negative |
| EK11 | Mycobacteria | EIEVDDDLIQK | -3.5 | |

**Supplementary Table 2. Hierarchy of peptide binding to HLA-E determined by peptide exchange ELISA assay screening**

Table of peptides predicted to bind HLA-E that were screened in the peptide exchange ELISA-based assay. Peptides are ranked in order of relative binding strength to HLA-E. Peptide ELISA signals were first normalised via background subtraction and subsequently expressed as a percentage of the positive control VL9 peptide. Peptide ID, origin and sequence are also reported. Peptides are subdivided into 11 categories based on relative HLA-E binding strength from those that generated signals below background levels to those that generated signals ranging from 70-100% of the positive control signal following background subtraction. Positive and negative controls were included in each ELISA plate to account for inter-plate variation. Biological repeats n=3, technical replica n=2.

#### Supplementary Figure 2

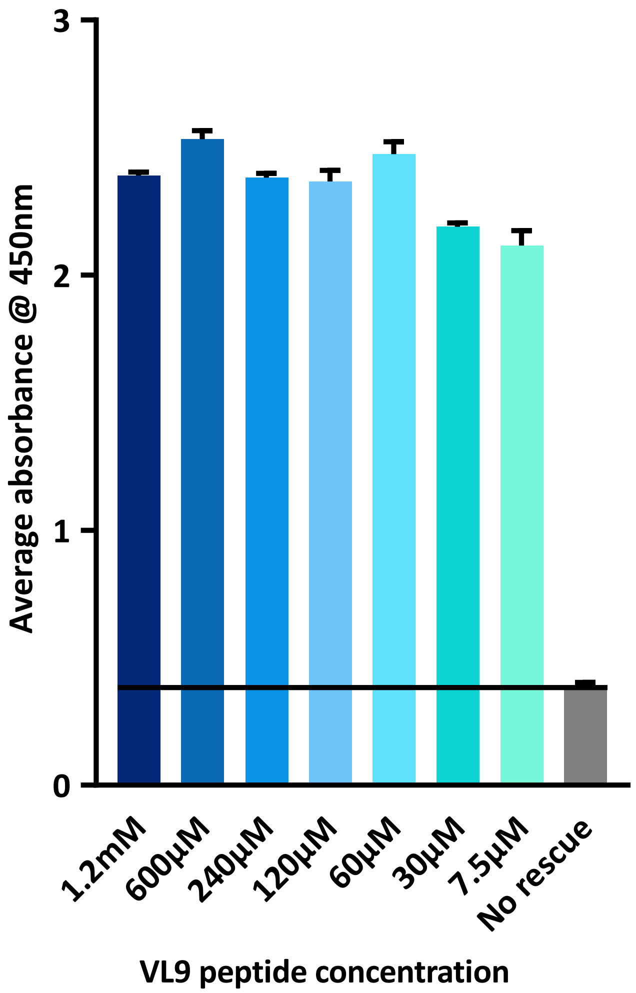


**Supplementary Figure 2. VL9 peptide titration in the optimised ELISA assay**
Average absorbance readings from a peptide exchange ELISA assay where the VL9 peptide was titrated from 1.2mM to 7.5µM. *Positive control peptide VL9 (HLA-B7-derived leader sequence peptide VMAPRTVLL) shown in dark blue with peptide free background (No rescue) in grey. Error bars depict standard error of the mean. Absorbance readings taken at 450nm. Biological repeats n=3, technical replica n=2. Stars above error bars reflect degree of significance in peptide binding by two-tailed t-tests: no star=P>0.05, *=P≤0.05, **=P≤0.01, ***=P≤0.001, ****P=≤0.0001.*

### Supplementary Table 3

| Peptide ID | Sequence | Organism | %VL9 rank | Structure | Ref. |
|---|---|---|---|---|---|
| Mtb44 | RLPAKAPLL | *Mycobacteria* | 96.5 | Yes | Walters, L,C. et al (2018) |
| IL9 | IMYNYPAML | *Mycobacteria* | 64.5 | Yes | Lucy C. Walters et al. unpublished |
| Mtb44 FP2 | RFPAKAPLL | *Mycobacteria primary anchor variant* | 57.3 | Yes | Walters, L,C. et al (2018) |
| Mtb44 FP9 | RLPAKAPLF | *Mycobacteria primary anchor variant* | 56.2 | Yes | Walters, L,C. et al (2018) |
| Mtb14 | RMAATAQVL | *Mycobacteria* | 48.4 | Yes | Lucy C. Walters et al. unpublished |
| RL9HIV | RMYSPTSIL | HIV | 43.4 | Yes | Walters, L,C. et al (2018) |
| BZLF1 | SQAPLPCVL | EBV | 40.7 | No* | N/A |
| Mtb44 QP2 | RQPAKAPLL | *Mycobacteria primary anchor variant* | 31.3 | Yes | Walters, L,C. et al (2018) |
| HIV Gag 9mer #6 | AISPRTLNA | HIV | 2.2 | No** | N/A |

**Supplementary Table 3. Table of peptide-bound HLA-E structures and corresponding ELISA binding scores**
Peptide-bound HLA-E crystal structures previously published by our lab in addition to unpublished structures and their corresponding rankings in the peptide exchange ELISA assay. Peptide binding scores are taken from supplementary figure 3 where inter-assay signals were normalised via background subtraction and subsequently expressed as percentages of the position VL9 peptide control. Mtb44 'FP2' = position 2 primary anchor substituted peptide where the wild type Leu was replaced with Phe. Mtb44 'FP9' = position 9 primary anchor substituted peptide where the wild type Leu was replaced for Phe. Mtb44 'QP2' = position 2 primary anchor substituted peptide where the wild type Leu was replaced for Gln.
*Despite an ELISA ranking above that of other crystallisable peptides (Mtb44 QP2), diffracting crystals were not obtained for HLA-E in complex with the EBV-derived BZLF1 peptide.
****Although crystallisation trials were set up for peptides with lower ELISA rankings, such as HIV Gag 9mer #6 (AISPRTLNA), crystals did not form. HIV Gag 9mer #6 is representative of a number of peptides that generated signals below 10% of VL9 following background subtraction and did not generate protein crystals.*
